# A CAMKK2-UBR4-19S Proteasome Axis Regulates Chondrocyte Proteostasis and SOX9 Stability

**DOI:** 10.64898/2026.05.07.723609

**Authors:** Xinchun Ding, Yong Li, Kasi Hansen, Amber L. Mosley, Elizabeth S. Yeh, Emma H. Doud, Uma Sankar

## Abstract

**Objective:** Investigate how Ca^2+^/calmodulin dependent protein kinase kinase 2 (CaMKK2) orchestrates a catabolic shift in chondrocytes during early osteoarthritis (OA).

**Methods:** Cartilage, osteochondral plugs and chondrocytes were collected from patients undergoing total hip arthroplasty or non-OA donors. SOX9 levels were assessed via immunoblotting or immunohistochemistry (IHC). Sox9 levels were also assessed by IHC in knee joints from wild-type (WT) and *Camkk2^-/-^* mice that underwent sham or destabilization of medial meniscus (DMM), with or without STO-609 (0.033 mg/kg) treatment. Co-immunoprecipitation followed by mass spectrometry was performed to identify CaMKK2 interacting proteins in chondrocytes. Kinase assays were analyzed by immunoblotting and phosphosites identified by mass spectrometry. Proteasome function was assessed in murine and human chondrocytes lacking or expressing kinase-active or kinase-inactive CaMKK2.

**Results:** Inhibition or loss of CaMKK2 increased SOX9, whereas the expression of kinase-active, not inactive, CaMKK2 reduced Sox9 in human and mouse OA cartilage. Proteomic analysis of CaMKK2 immunoprecipitates revealed the presence of ubiquitin E3 ligase Ubr4 and the 19S proteasome regulatory particle (RP). CaMKK2 kinase activity was dispensable for its interactions with Ubr4, 19S RP, and Sox9–ubiquitin conjugates, and kinase-inactive CaMKK2 attenuated Sox9 degradation in chondrocytes. Further, CaMKK2 phosphorylated the 19S RP ATPase Psmc5 on Ser136, and an intact kinase increased proteasome activity in chondrocytes.

**Conclusions:** Our findings identify CaMKK2 as a dual-function regulator of chondrocyte UPS with a scaffolding role to assemble UPSUbr4-19S RP around polyubiquitinated proteins such as Sox9, and a catalytic role to enhance proteasome function, potentially through Psmc5 phosphorylation, thereby linking chondrocyte inflammatory signaling to Sox9 degradation and cartilage degeneration.

## Introduction

Despite its prevalence and impact [1], no FDA-approved disease-modifying OA drugs (DMOADs) exist. A major obstacle to therapeutic development is the limited understanding of early molecular events driving OA. Cartilage injury elicits an immediate inflammatory response in the chondrocytes through the release of proinflammatory cytokines such as interleukin (IL)-1β and tumor necrosis factor (TNF)-α [2]. These cytokines promote extracellular Ca^2+^ influx into chondrocytes via ion channels such as transient receptor potential vanilloid 4 (TRPV4). This transiently increases intracellular Ca^2+^, which amplifies inflammation and stimulates cartilage matrix degradation by matrix metalloproteinases (MMPs) and aggrecanases [3–5].

Mechanical injury to the joint also triggers a rapid increase in polyubiquitination of target proteins and induces a distinct ubiquitin signature in chondrocytes [6]. The ubiquitin-proteasome system (UPS), a Ca²⁺-dependent protein degradation machinery, is activated following joint injury and it promotes inflammation by accelerating the degradation of IκB kinase (IκK) and leading to the activation of nuclear factor (NF)-κB, a key transcription factor that drives inflammatory gene expression in chondrocytes [6–8]. This interplay between inflammation, Ca^2+^ signaling, and UPS activity shifts chondrocytes to a proinflammatory, catabolic state by degrading proteins involved in cartilage synthesis and maintenance and preserving those promoting chondrocyte hypertrophy and death, and cartilage degradation [6, 9].

SRY-box 9 (Sox9), a master transcription factor regulator of chondrocyte differentiation and a direct activator of collagen type II alpha 1 (Col2a1) and aggrecan, is rapidly degraded through the UPS in response to IL-1β, TNF-α, and NF-κB activation [10–12]. Notably, proteasome inhibitors such as Bortezomib and MG-132 protect against inflammation and cartilage degradation in human chondrocytes and rodent PTOA models [13–16]. However, the regulation of UPS in chondrocytes and its role in OA pathogenesis remain poorly understood.

We previously demonstrated that CaMKK2, a Ser/Thr kinase activated by intracellular Ca^2+^ transients, is elevated in human and murine OA articular chondrocytes [17, 18]. Inhibition or ablation of CaMKK2 protected against PTOA in mice by inhibiting IL-6/signal transducer and activator of transcription 3 (STAT3)/MMP13 pathway mediated inflammatory, catabolic and apoptotic responses downstream of IL-1β in chondrocytes [17, 18]. CaMKK2 deletion counteracted the anti-anabolic effects of IL-1β in chondrocytes by increasing *Col2a1* mRNA and preserving Col2 and aggrecan protein levels [18].

Given that Sox9 is the upstream regulator of *Col2a1* [10, 19], we hypothesized that CaMKK2 played a role in maintaining Sox9 levels in inflamed chondrocytes. Our studies suggest that CaMKK2 stabilizes UPS in chondrocytes, thereby promoting catabolic pathways that drive OA pathogenesis.

## Materials and Methods

### Human sample procurement and preparation

Human OA tissue collection was approved by Indiana University Institutional Review Board (IRB ID 2003626223) and Department of Defense Human Research Protection Office (HRPO E01450.1a). De-identified human femoral heads were collected from patients undergoing total hip arthroplasty (THA) for primary OA as part of a previous study [17]. Briefly, radiography-based Kellgren-Lawrence (KL) grade 3 or lower was used as a cut off [20]. Femoral heads (N=6 female; **Fig. 1-Table 1**) were immediately placed in sterile saline prior to processing as described before [17]. Cartilage was shaved from OA and intact portions of the same femoral head, cut into 1×1×0.5 mm sections, lysed and processed for immunoblotting. For OA explant culture, osteochondral cores (2 cores each from 4 individual patients) were extracted using an OATS instrument (AR-1981-10H, Arthrex Inc., Naples, FL). OA severity of the specimens was histologically determined and reported before [17].

**Figure 1:**
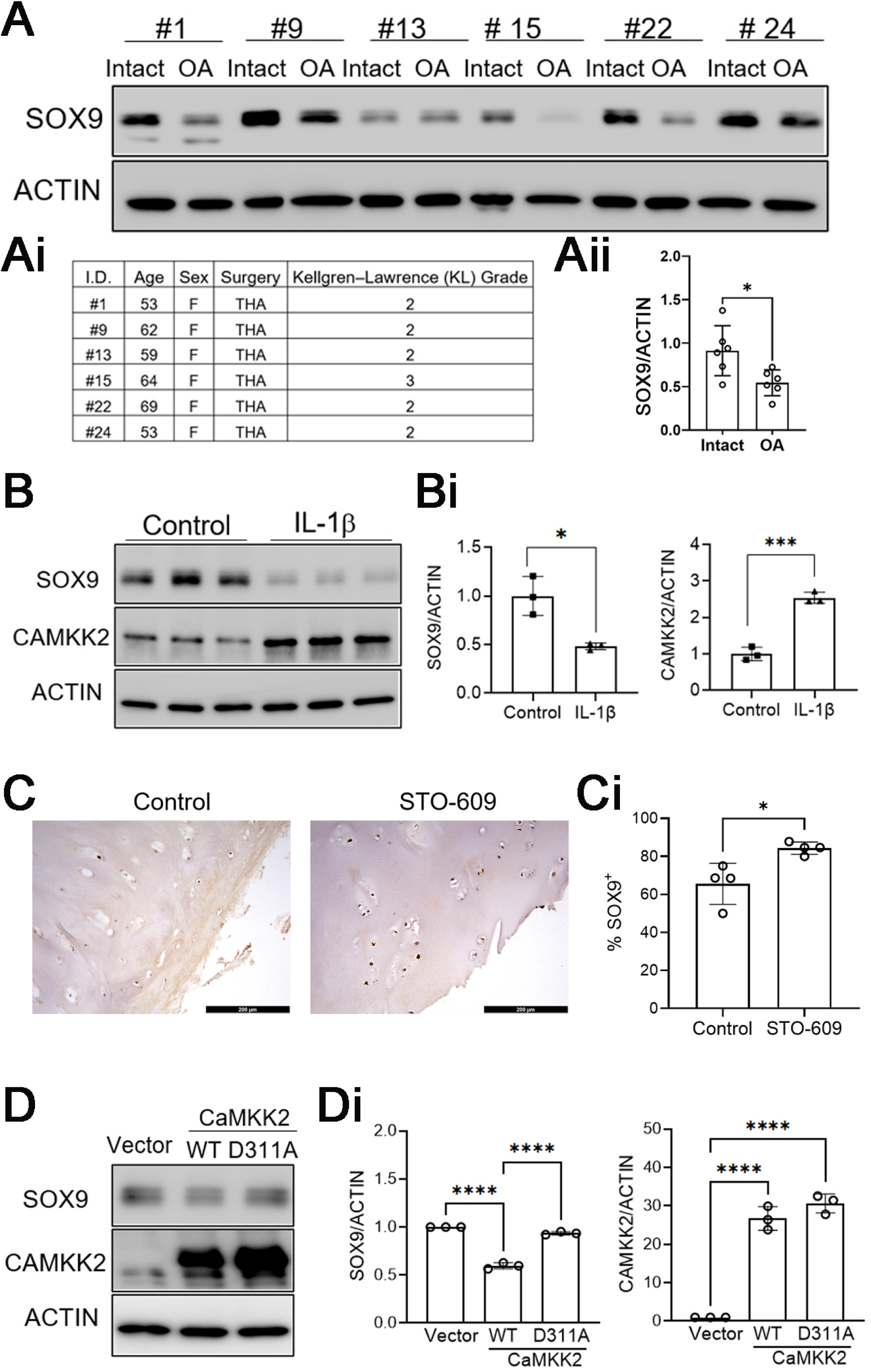
CAMKK2 inhibition elevates SOX9 in human OA cartilage. **(A)** Immunoblots depicting SOX9 and ACTIN (loading control) protein levels in human cartilage isolated from “intact” versus OA portions of the same femoral head of individual THA patients (n=6 pairs). **(Ai)** Table shows the patient demographic details for the de-identified human THA OA cartilage samples used in this study. **(Aii)** Quantification of immunoblots from 1A. Error bars = standard deviation (SD); *p-*values: ** *p*<0.01. **(B)** Immunoblots from P0 chondrocytes, isolated from healthy human donors (NDRI), treated with 1X PBS (Control) or 10 ng/ml IL-1β for 24 h probed for CAMKK2 and ACTIN (loading control) immunoreactivity. Each lane represents lysates from an individual donor, indicating N=3 biological replicates. **(Bi)** Ratio of average signal intensities of SOX9 to ACTIN and CAMKK2 to ACTIN. SD; *p-*values: * *p*<0.05; *** *p*<0.001. **(C)** Representative IHC images (200X; scale bar=100 µm; n=4/group) of human THA OA cartilage explants treated with 1X PBS (Control) or 8 µM STO-609 for 24 h, showing immunoreactivity to SOX9. **(Ci)** Quantification of SOX9 positivity in control and STO-609 treated OA cartilage explants (n=4 individual OA explants/cohort). SD; *p-*values: * *p*<0.05. **(D)** Immunoblots of cell extracts from healthy human chondrocytes, isolated from NDRI donors, infected with Lentivirus vector (vector) or Lentiviruses expressing rat CaMKK2 (WT) or a kinase-inactive CaMKK2 mutant (D311A) were probed for CAMKK2, SOX9 and ACTIN. **(Di)** Average signal intensities of SOX9 and CAMKK2, compared to ACTIN from n=3 independent experiments (biological replicates) are shown. SD; ****p < 0.0001.

**Table 1.** Patient demographic details for de-identified human THA OA cartilage samples used for explant cultures and IHC:

| ID | Age | Sex | Surgery | Kellgren–Lawrence (KL) Grade |
| --- | --- | --- | --- | --- |
| 1 | 57 | F | THA | 2 |
| 3 | 57 | M | THA | 2 |
| 4 | 70 | M | THA | 3 |
| 5 | 43 | M | THA | 2 |

### Human OA explant culture

OA cores/explants were immediately rinsed in 1X PBS and placed in culture media for 24 h. Explants from the same patient were “paired” and treated for 48 h with either 1× PBS (Control) or STO-609 (8 µM; TOCRIS, MO, USA), fixed in 4% paraformaldehyde (PFA) for 48 h, decalcified in 14% EDTA (pH 7.4), paraffin-embedded and cut into 5 μm thick sections. STO-609 was prepared as before [18].

### Mice

Mouse studies were approved by Indiana University School of Medicine (IUSM) Institutional Animal Care and Use Committee (IACUC), and all experiments were performed in compliance with NIH guidelines on the use and care of laboratory and experimental animals. WT and *Camkk2^−/−^* mice [18] (both C57BL/6) were housed in the IUSM Laboratory Animal Resource Center under 12 h light/dark cycle. Food and water were provided *ad libitum*.

### Sham/DMM Surgeries and sample collection

Sham and destabilization of the medial meniscus (DMM) surgeries were originally performed on 10-week-old WT (n=12) and *Camkk2^−/−^* mice (n=12) as part of a previous study [18]. Half of the DMM-WT mice received intraperitoneal (i.p.) injections of CaMKK2 inhibitor STO-609 (0.033 mg/kg) thrice a week (w), starting from the day after the surgery until termination at 12 w post-surgery, while the other groups received tri-weekly i.p, injections of sterile saline. Articulated knee joints were then isolated, fixed in 4% PFA, decalcified and processed for histology or IHC. OA severity, synovial inflammation and subchondral bone sclerosis from these mice were reported previously [18]. For this study, we analyzed n=3 samples/group for Sox9 IHC.

### IHC of human explant and mouse knee joint sections

Human explant and/or mouse knee joint sections were blocked with nonfat BSA (1% in 1 x PBS) for 1 h and then incubated with αSox9 (Sigma; **Table 2**) at 4C overnight. Tissue sections were washed, incubated with biotinylated secondary antibody for 1 h at RT, and counterstained with Gill No. 1 Hematoxylin (Millipore Sigma). Signals were amplified using a Vectastain Elite ABC-HRP kit (Vector Laboratories, Burlingame, CA). Images were captured using a Leica DMi8 microscope, processed with Leica LAS-X software (Leica CM1950, Wetzlar, Germany), and quantified by counting immune-positive and total cells within articular cartilage from 7 regions of interest per sample using ImageJ (NIH, USA) as before [17].

**Table 2.** List of primary antibodies used and their application in this study:

| Primary Antibody | Vendor | Catalog Number | Antibody Dilution | Application |
| --- | --- | --- | --- | --- |
| Rabbit αCaMKK2-NT | US Biological, Salem, MA | 033168 | 1:250 | IP |
| Mouse αCaMKK (pan) | BD, Franklin Lakes, NJ | 610544 | 1:1000 | IB |
| Rabbit α-pCaMKIV (T196, T200) | Invitrogen, Waltham, MA | PA5-105011 | 1:1000 | IB |
| Mouse αCaMKIV | BD | 610275 | 1:1000 | IB |
| Rabbit αPhospho-S/T | Abcam | Ab17464 | 1:1000 | IB |
| Rabbit αSox9 | Sigma, St. Louis, MO | AB5535 | 1:1000 | IB, IP, IF, IHC |
| Rabbit anti-UBR4 | Abcam, Waltham, MA | Ab86738 | 1:1000 | IB |
| Rabbit anti-Psmd2 | Abcam | Ab140675 | 1:1000 | IB |
| mouse anti-Ubiquitin | Cell signaling, Danvers, MA | 3936 | 1:1000 | IB |
| mouse anti-Pmsc5 | Santa Cruz, Dallas, TX | Sc-390631 | 1:1000 | IB, IP |
| Mouse anti-b-Actin | Sigma | A1978 | 1:10,000 | IB |
| IHC – Immunohistochemistry; IB – Immunoblotting; IF – Immunofluorescence; IP – Immunoprecipitation |  |  |  |  |

### Primary human articular chondrocyte isolation and culture

Human chondrocytes were isolated from non-OA donor femurs procured from National Disease Research Interchange (NDRI) and chondrocytes isolated as previously described [17]. P0 chondrocytes were serum starved overnight, released into media containing 10% FBS and treated with 10 ng/ml human recombinant IL-1β (Millipore Sigma) as indicated.

### Primary murine articular chondrocyte (MAC) isolation and culture

MACs were isolated from knee joints of 4–6-day-old *WT* and *Camkk2^−/−^* mice; cartilage, devoid of epiphyseal tissues, was pooled from 6 to 8 pups per genotype, minced and sequentially digested with two different concentrations of collagenase II (Millipore Sigma) and plated as previously described [18]. P0 chondrocytes were serum starved for 12 h, released into full media before treating with 10 ng/ml IL-1β, with or without 1 µM MG132 (Cayman Chemical, Ann Arbor, MI).

### Immunofluorescence staining

MACs were fixed for 15 min in 4% PFA, and permeabilized for 10 min in 0.02% TritonX-100. Cells were then blocked with 5% normal goat serum for 1 h, incubated in rabbit-αSox9 at 4C overnight and fluorescence-conjugated secondary antibody for 1 h at RT followed by co-staining with DAPI.

### Lentivirus vector production and transduction into primary chondrocytes

The cloning of FLAG-rat (r) CaMKK2 (kinase-active; WT) and/or FLAG-rCaMKK2 D311A (kinase-inactive) cloned into the dual-promoter Lentivirus vector pCDH-MSCV-MCS-EF1-copGFP (System Biosciences (SBI), Mountain View, CA), generation of Lentivirus particles, and determination of virus titers have been previously described [17]. Lentiviruses (control vector, WT or mutant CaMKK2) were added at an MOI of 5:1 in the presence of the cationic polymer hexadimethrine bromide (polybrene; 4 μg/ml; Millipore Sigma) on P0 chondrocytes and cultured for 7 days before preparing protein lysates.

### Immunoblotting

Equal amount of protein from cartilage, MACs or human chondrocytes was fractionated under denaturing conditions on SDS-PAGE and transferred onto Immobilon-P membranes (Millipore Sigma). Blocking, primary, and secondary antibody incubations were performed in Tris-buffered saline with Tween-20 (TBST) containing 5% non-fat dry milk. Membranes were probed with primary antibodies **(Table 2)** and horseradish peroxidase-conjugated secondary antibodies (Jackson Immunoresearch Laboratories, West Grove, PA, USA). The target proteins were visualized with ChemiDox MP Image System and band densities quantified by using Image Lab (all Bio-Rad).

### Co-Immunoprecipitation

Following indicated treatments, primary MACs were placed into iced tween lysis buffer (25 mM Hepes (pH 7.5), 50 mM NaCl, 25 mM NaH2PO4, 0.5% Tween 20, 10% glycerol, 1 mM dTT, 10 mM β-glycerophosphate, 2 mM EGTA, 2 mM EDTA, 25 mM NaF, 1 mM sodium vanadate, 1 mM PMSF, 1 mg/ml aprotinin, 1 mg/ml leupeptin, 10 mg/ml pefabloc, and 100 nM okadaic acid), sonicated on ice, centrifuged 14000 rpm for 30 min at 4^0^C. About 500 µg total protein was precleared with 5 µl of protein G mag Sepharose (Millipore Sigma) and incubated with 5 µg rabbit IgG or rabbit-αCaMKK2-NT antibody (US Biological) or rabbit-αSox9 and 5 µl protein G mag Sepharose beads for 4h at 4^0^C. The beads were spun down for downstream analyses using immunoblotting or mass spectrometry.

### Kinase Assay

His-tagged recombinant human PSMC5 (0.2 µg; Ag6549, Proteintech, Rosemont, IL) and GST-tagged recombinant human CAMKK2 (0.1 µg; 40046, BPS Bioscience, San Diego, CA) were mixed in 20 µl of kinase buffer (50 mM HEPES, pH 7.5; 300 mM NaCl; 10 mM MgCl₂; 400 µM ATP; 5 µM calmodulin; 1 mM CaCl₂; 1 mM DTT) and incubated at 30 °C for 30 min with gentle shaking. PSMC5 (0.2 µg) incubated in kinase buffer alone served as negative control. For positive control, recombinant human CAMKIV (5 µg/ml; NBP1-78878, Novus Biologicals, Centennial, CO) was incubated with CAMKK2 (0.1 µg) in kinase buffer. Two µl were resolved by SDS-PAGE and immunoblotted to assess protein phosphorylation. The remaining kinase reaction was analyzed by mass spectrometry (LC-MS/MS).

### Mass Spectrometry

Sample preparation, mass spectrometry analysis, bioinformatics, and data evaluation for quantitative proteomics experiments were performed in collaboration with the Indiana University School of Medicine Center for Proteome Analysis similarly to several previously published protocols.[21, 22]

### On bead digests

After washing, beads were covered with 8 M Urea, 50mM Tris hydrochloride, pH 8.5, reduced with 5mM tris (2-carboxyethyl) phosphine hydrochloride (TCEP, Sigma-Aldrich Cat No: C4706) for 30 minutes at room temperature to reduce the disulfide bonds. The resulting free cysteine thiols were alkylated using 10 mM chloroacetamide (CAA, Sigma Aldrich Cat No: C0267) for 30 minutes at RT, protected from light. Samples were diluted to 2 M Urea with 50 mM Tris pH 8.5 and proteolytic digestion was carried out with Trypsin/LysC Gold (0.5 µg, Mass Spectrometry grade, Promega Corporation Cat No: V5072) overnight at 35 °C. After digestion, samples were desalted on Pierce C18 spin columns (Thermo Fisher Scientific Cat No:89870) with a wash of 0.5% TFA followed by elution in 70% acetonitrile 0.1% formic acid (FA). Samples were dried down by speed vacuum and resuspended in 20uL 0.1% formic acid.

#### LC-MS/MS

Approximately 1/5^th^ of each sample was then injected onto a 25 cm MicrOmics C18 column (PN:HC1A-JFST, MicrOmics technologies) using an EasyNano1200 LC coupled to an Exploris 480 orbitrap mass spectrometer (Thermo Fisher Scientific). Solvent B (80% Acetonitrile, 0.1 % FA) was increased from 8-35 % over 90 minutes, increased from 35-65 % over 15 min, increased to 85% over 5 min, held at 85% for 5 min, and decreased to 4% over 5 min all at 300 nL/min. The mass spectrometer was operated in positive ion mode, advanced peak determination on, default charge state of 2 and no user defined lock mass. 3 FAIMS CVs (-40, -55, and -70) were used with identical MS1 and MS2 parameters. 1.3 second cycle time was used with MS1 parameters of scan range 375-1500 m/z: orbitrap resolution of 60,000, standard AGC, automatic max IT, and RF lens of 40%. Monoisotopic peak determination was set to peptide with a minimum intensity filter of 5.0e3, charge state filter of 2-7, and dynamic exclusion of 30 s with 5 ppm mass tolerance. 1.3 sec top S cycle time was used for fragmentation. MS2 parameters included an isolation window of 1.6 m/z, normalized high energy dissociation energy of 30 %, orbitrap resolution of 15,000, user defined first mass of 110 m/z, standard AGC target and auto max IT. Data were recorded using Tune application 4.2.362.42.

### Data analysis

Resulting RAW files were analyzed in Proteome Discover™ 2.5.0.400 (Thermo Fisher Scientific) searched with *Homo sapiens* UniProt reviewed proteome FASTA (downloaded 051322, 20272 sequences) and common laboratory contaminants (73 sequences) (for recombinant kinase assay results additional protein databases included were: *E. coli* UniProt FASTA (downloaded-110222, 4305 sequences), SF9 reviewed FASTA (49 sequences)) [23]. SEQUEST HT searches were conducted with full trypsin digest, 3 maximum number missed cleavages; precursor mass tolerance of 10 ppm; and a fragment mass tolerance of 0.02 Da. Static modifications used for the search were: 1). carbamidomethylation on cysteine (C) residues. Dynamic modifications used for the search were 1) oxidation on M 2) phosphorylation on S, T or Y, and 3) Acetylation on K. A maximum of 3 dynamic modifications were allowed per peptide. Percolator false discovery rate (FDR) filtration of 1% was applied to both the peptide-spectrum match and protein levels. IMP-ptmRS node was used for phosphosite localization scoring.

### Proteasome Assay

Proteasome activity in MACs and human chondrocytes was measured using Proteasome Assay kit (Cayman chemicals; cat # 10008041) per manufacturer’s protocol. Briefly, cells were seeded in 6-well plates (5X10^5^ MACs or 1X10^6^ human chondrocytes per well) and allowed to adhere overnight, before treating IL-1β for 24 h or Lentiviruses for 7 days. Cell lysates were then prepared and transferred to opaque 96-well plates (Sigma). The proteasome substrate, SUC-LLVY-AMC was added to the wells and fluorescence intensities were measured using a FLx-800 fluorimeter (Bio-Tek) with the absorption (excitation) set at 360 nm and the emission set at 480 nm.

### Statistical analyses

Statistical analyses were performed using GraphPad Prism (GraphPad, San Diego, CA). Normality assumptions were evaluated using histograms and QQ plots. For normally distributed data, Student’s paired *t*-test (2 sample comparisons) or ordinary one-way ANOVA followed by Tukey’s HSD post-hoc (comparing >2 groups) were used. For non-normal data, non-parametric tests: Mann–Whitney test for 2 sample comparisons or Kruskal–Wallis test followed by Dunn’s post hoc to compare >2 groups were used. Significance was set at *p*<0.05. Data are represented as mean ± standard deviation (SD).

## Results

### CAMKK2 regulates SOX9 in human OA chondrocytes

SOX9, the master regulator of chondrocyte degradation, maintains cartilage homeostasis by directly regulating the expression of cartilage extracellular matrix (ECM) components such as COL2 and aggrecan. Its inactivation accelerates cartilage degradation and promotes OA [10, 24–26]. *SOX9* mRNA is reported to be downregulated in human OA cartilage [27, 28], but the fate of SOX9 protein in human OA has not been reported. To assess SOX9 protein levels, we analyzed hip OA cartilage from female patients undergoing THA (KL grade 2–3). Compared to intact regions of the same femoral head, SOX9 levels were reduced by 1.8-fold in OA cartilage (n = 6 pairs, **Fig. 1A-Aii**). Primary non-OA human articular chondrocytes exposed to IL-1β (10 ng/mL, 24 h) displayed a 2-fold decrease in SOX9 **(Fig. 1B-Bi)**. In contrast, IL-1β increased CAMKK2 by 2.5-fold in human chondrocytes as reported [17]. Treatment of OA cartilage explants with CaMKK2 inhibitor STO-609 [17, 18, 29], significantly increased SOX9 levels, implicating CAMKK2 in the regulation of SOX9 (**Fig. 1C-Ci**).

Lentivirus-mediated expression of kinase-active CAMKK2 in human chondrocytes reduced SOX9 by 1.8-fold, whereas that of kinase-inactive CaMKK2 had no effect **(Fig. 1D-Di),** indicating that CaMKK2 kinase activity regulates SOX9 levels.

### Regulation of Sox9 by CaMKK2 in mouse articular chondrocytes

Sox9 is downregulated in the mouse articular cartilage following the destabilization of medial meniscus (DMM), whereas CaMKK2 is upregulated [18, 30]. To test whether inhibiting or ablating CaMKK2 will preserve Sox9 in murine DMM cartilage, we performed sham or DMM surgeries in 10-week (w) -old male wild-type (WT) and *Camkk2^-/-^* mice and treated half of the DMM-WT mice with STO-609. Sox9 positivity in articular cartilage was significantly reduced in WT mice 12 w post-DMM, whereas treatment with STO-609 prevented Sox9 downregulation. Notably, Sox9 levels remained unchanged in *Camkk2^-/-^* articular cartilage mice following DMM **(Fig. 2A-Ai)**.

**Figure 2:**
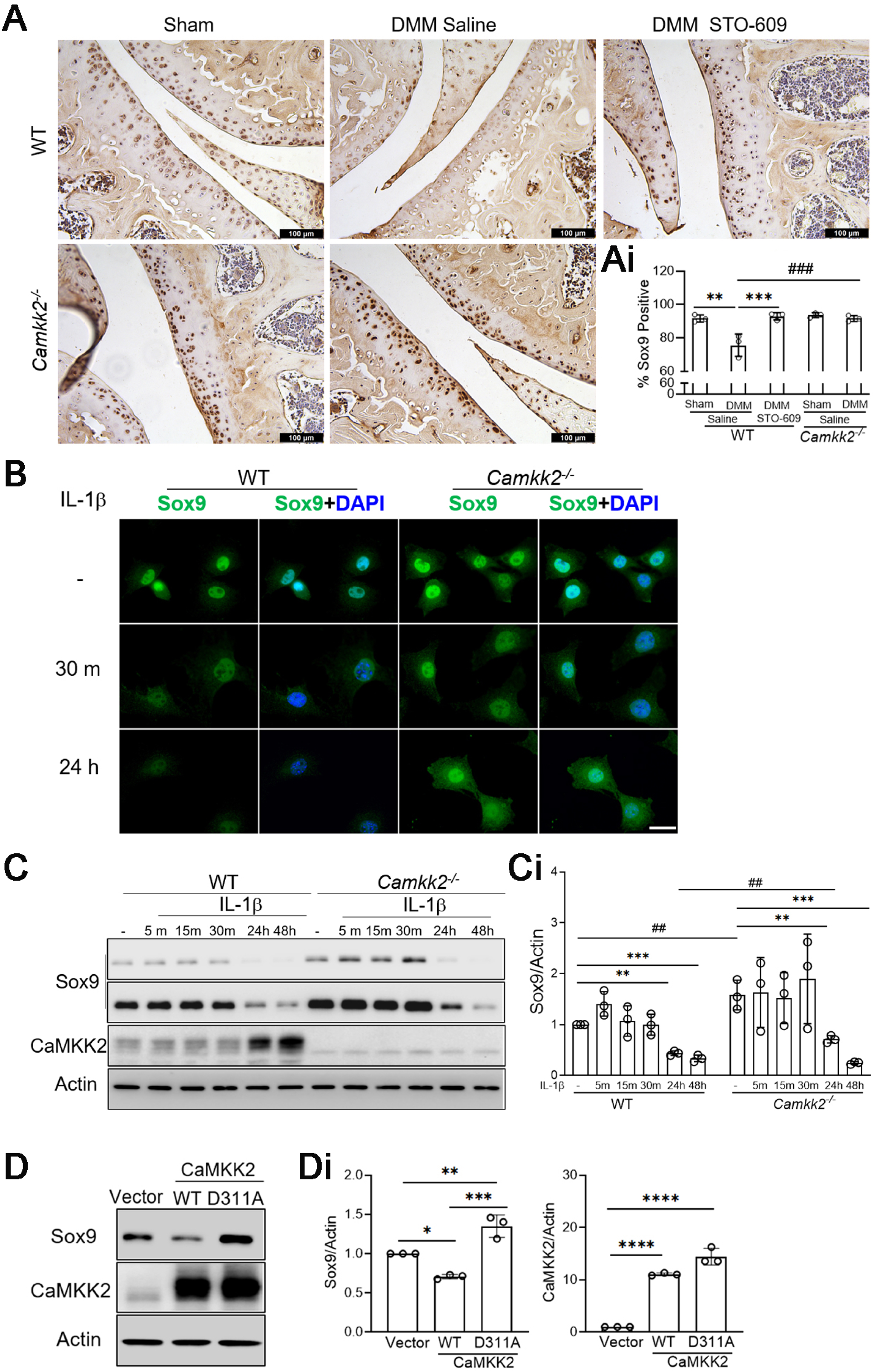
Absence or loss of CaMKK2 elevates Sox9 and protects against its loss in inflamed murine cartilage. **(A)** Representative IHC images (200X) of articular cartilage sections from WT-sham, WT-DMM-saline, WT-DMM-STO-609, *Camkk2^-/-^*-sham or *Camkk2^-/-^*-DMM mice collected at 12 w post-surgery and probed for Sox9 immunoreactivity. **(Bottom Right)** Quantification of Sox9 IHC in indicated genotypes (n=3/cohort). SD; *p*-values: ** *p* < 0.01 and \*\*\**p* < 0.001 compared to WT-sham; ^###^ *p*<0.001 compared to WT-DMM-saline. **(B)** Immunofluorescence images (representative of n=3 biological replicates/group) depicting Sox9 (Green) levels and DAPI nuclear staining in primary WT and *Camkk2^-/-^* MACs treated without or with 10 ng/ml IL-1β for 30 min or 24 h. **(C)** Immunoblots of cell extracts from WT and *Camkk2^-/-^* iMACs untreated (-) or treated with 10 ng/ml of IL-1β for indicated time points and probed for Sox9, CaMKK2 and b-Actin immunoreactivity. The two Sox9 blots represent lighter and darker exposures of the same blot. Representative blots from n=3 independent experiments are shown. **(Ci)** Average signal intensities of Sox9 normalized to those of Actin from n=3 blots. SD; * indicates *p*-values for within genotype comparison and # for WT vs. *Camkk2^-/-^*comparisons; ** *p* < 0.01 and \*\*\**p* < 0.001; ^##^ *p*<0.01. **(D)** Immunoblots of primary WT MACs, infected with Lentivirus vector only (vector) or expressing CaMKK2 (WT) or CaMKK2 mutant (D311A) and probed for CaMKK2, Sox9 and Actin are shown. **(Di)** Average signal intensities of Sox9 and CaMKK2, compared to Actin from n=3 independent experiments are shown. SD; * *p*<0.05; ** *p* < 0.01; \*\*\**p* < 0.001 and ****p < 0.0001.

Immunofluorescence confirmed higher Sox9 in naïve *Camkk2^-/-^* primary mouse articular chondrocytes (MACs) compared to WT. Within 30 min of IL-1β treatment, Sox9 sharply declined in WT cells but persisted at higher levels in *Camkk2^-/-^* MACs **(Fig. 2B)**. At 24 h of IL-1β treatment, *Camkk2^-/-^* MACs showed increased cytoplasmic Sox9, while retaining substantially higher nuclear and total Sox 9 levels than WT **(Fig. 2B)**.

Next, we examined Sox9 levels in IL-1β-treated WT and *Camkk2^-/-^*MACs and found it to be reduced by 2.5-fold within 24 h and by 3-fold within 48 h in WT MACs **(Fig. 2C-Ci)**. *Camkk2^-/-^* MACs exhibited higher basal Sox9 levels compared to WT (1.6-fold) and maintained elevated Sox9 during the first 30 min of IL-1β exposure (1.3–1.9-fold) **(Fig. 2C-Ci)**. Sox9 in *Camkk2^-/-^* MACs decreased by 2.2-fold at 24 h of IL-1β treatment, but it remained 1.7-fold higher than WT. By 48 h of IL-1β treatment, Sox9 levels were similar in both genotypes **(Fig. 2C-Ci)**. Expression of kinase-active CaMKK2 reduced Sox9 by 1.4-fold in WT MACs, whereas that of kinase-inactive CaMKK2 increased it **(Fig. 2D-Di)**. Thus, loss of CaMKK2 or inhibition of its activity elevates Sox9 in chondrocytes and protects against its rapid depletion due to inflammation.

*Sox9* mRNA levels were comparable between naïve WT and *Camkk2^-/-^* MACs **(Supplementary Fig. 1),** suggesting that the regulation by CaMKK2 is transcriptional. Sox9 protein levels are higher in CaMKK2 null MACs, but IL-1β treatment diminishes its levels in both WT and *Camkk2^-/-^* MACs. To understand the involvement of the proteasome in triggering IL-1β-mediated Sox9 degradation in WT and *Camkk2^-/-^* chondrocytes, we blocked proteasome activity using 1 µM MG-132 [31]. Sox9 levels were significantly elevated at baseline in *Camkk2^-/-^* MACs and remained higher at 4 h post IL-1β treatment, compared to WT **(Supplementary Fig. 2)**. MG-132 protected Sox9 from IL-1β-mediated degradation in WT and *Camkk2^-/-^* MACs, indicating proteasome involvement in Sox9 turnover in both genotypes.

### CaMKK2 interacts with components of the chondrocyte ubiquitin proteasome system

To elucidate the mechanism by which CaMKK2 regulates Sox9 levels, we performed co-immunoprecipitation (co-IP) of CaMKK2 from IL-1β-treated WT MACs followed by liquid chromatography–tandem mass spectrometry (LC-MS/MS) to identify associated proteins **(Fig. 3A)**. This time point was selected because CaMKK2 levels increased by 7.5 fold in MACs 24 h after IL-1β treatment [18]. Successful enrichment of CaMKK2 in the immunoprecipitates indicated efficient capture of interacting proteins by αCaMKK2 antibody **(Fig. 3B)**. Ubiquitin protein ligase E3 component N-Recognin 4 (Ubr4), a conserved E3/E4 ubiquitin ligase [32–34], was the most abundant protein (135 spectral counts) bound to CaMKK2 **(Fig. 3B)**. The large Ubr4 complex (≥600 kDa) includes its cofactors: calmodulin and potassium channel modulatory factor 1 (Kcmf1), an E3 ubiquitin ligase,[35] and both co-IP’d with CaMKK2 **(Fig. 3B-C)**.

**Figure 3:**
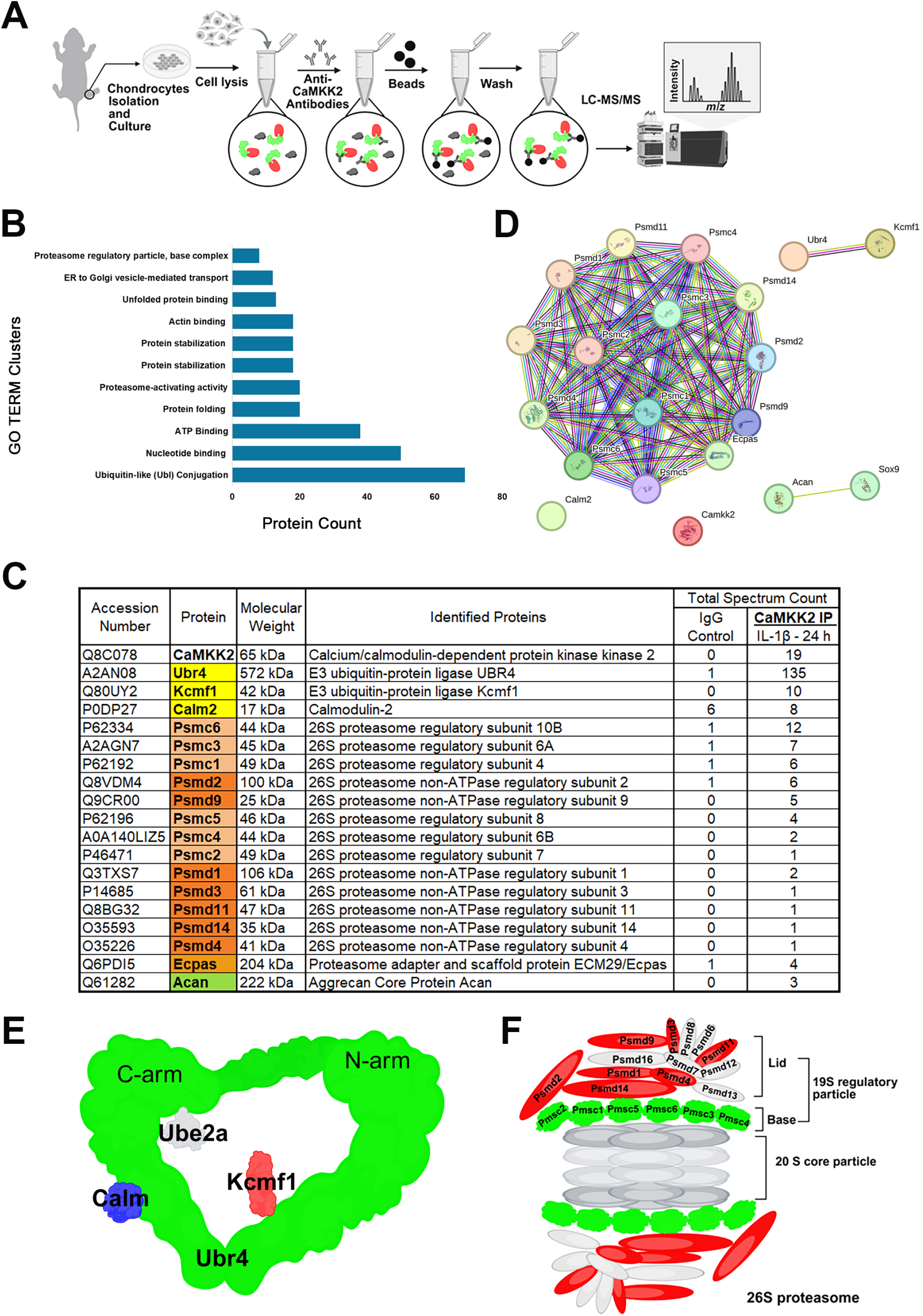
CaMKK2 interacts with the UPS in chondrocytes. **(A)** Schematic overview of the experimental workflow. Cell lysates prepared from primary WT MACs treated with 10 ng/ml IL-1β for 24 h were immunoprecipitated with control IgG or αCaMKK2 antibody followed by LC-MS/MS analysis to identify the interacting proteins. **(B)** Graph showing Gene Ontology (GO) Term functional annotation clustering of 241 proteins bound to αCaMKK2 antibody at ≥2-fold than IgG controls; generated from DAVID (Database for Annotation, Visualization and Integrated Discovery). **(C)** Table showing a list of CaMKK2 binding proteins that includes ubiquitin E3 ligases Ubr4 and Kcmf1, 19S proteasome RP subunits, Ecpas and Acan. Total spectrum counts in IgG control IP and CaMKK2 IP samples are shown. Color coding indicates protein complexes:– Yellow – Ubr4, Kcmf1 and Calm2 (Ubr4 complex); Light Orange – Psmc1-6 (19S RP base complex); Dark Orange – Psmd1-4, 9, 11, 14 (part of 19S RP lid complex); Light Brown – Ecpas, and Green – Acan. **(D)** STRING analysis of the CaMKK2 interacting proteins shown in **C – Table** plus Sox9, showing physical and functional associations. **(E)** Depiction of Ubr4 ring complex containing Ubr4, Kcmf1 and Calm, all of which bind to CaMKK2. **(F)** The 26S proteasome complex depicting components of the 19S RP lid and base subcomplexes. A, E and F were created with BioRender.com.

The 19S regulatory particle (RP) of the 26S proteasome was the other major interacting partner of CaMKK2 (**Fig. 3B)**. The 26S proteasome is composed of two subcomplexes: the 19S RP and the 20S catalytic core. The 19S RP is responsible for the recognition of polyubiquitinated substrates, their unfolding, deubiquitylation, and translocation into the 20S catalytic core which performs the proteolysis [36]. The 19S RP consists of a lid containing 16 ATP-independent subunits (PSMD1–16) and a base composed of six ATP-dependent subunits (PSMC1–6) **(Fig. 3D)**. Our proteomics data revealed CaMKK2 complexing with all six PSMC ATPases as well as PSMD1, PSMD2, PSMD3, PSMD4, PSMD9, PSMD11, and PSMD14 in chondrocytes **(Fig. 3B)**. Among these, PSMC6 showed the highest enrichment, followed by PSMC3, PSMC1, PSMD2, PSMD9, and PSMC5 **(Fig. 3B)**. The proteasome adapter/scaffold protein ECM29/Ecpas and aggrecan (Acan) were also detected in the CaMKK2 co-IP complex (**Fig. 3B)**. Collectively, these findings show that CaMKK2 interacts with key components of the chondrocyte UPS.

To validate the IP-proteomics findings, we treated primary WT MACs with 10 ng/mL IL-1β for 1 h or 24 h; co-IP’d cell lysates with αCaMKK2 and/or αSox9 antibodies and performed immunoblotting **(Fig. 4A)**. Co-IP confirmed CaMKK2 interaction with Ubr4 (∼600 kDa) and Psmd2 (∼100 kDa) in both naïve and IL-1β-treated MACs **(Fig. 4B)**. Sox9 co-IP analysis revealed the presence of Ubr4 and Psmd2, with IL-1β treatment enhancing their interaction with Sox9 **(Fig. 4C)**. Notably, IL-1β treatment did not alter total Ubr4 or Psmd2 levels in WT MACs; however, it increased CaMKK2 and reduced Sox9 levels **(Fig. 4B-F, Input)**. These data validate the LC-MS/MS data and suggest that IL-1β strengthens the association of Sox9 with Ubr4 and Psmd2, two key UPS components.

**Figure 4:**
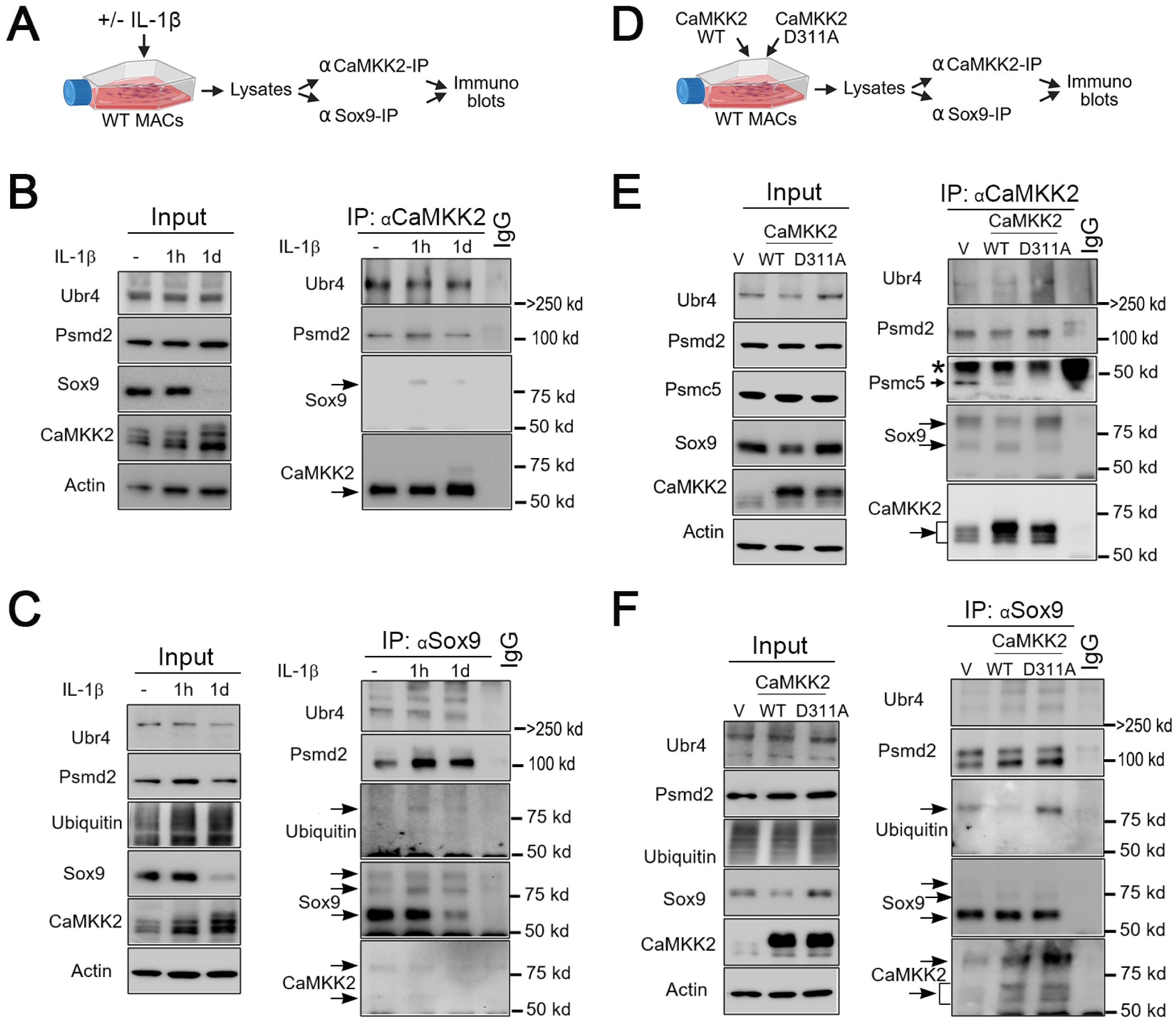
Association of CaMKK2 with Ubr4, Psmd2, Psmc5 and polyubiquitinated Sox9 in chondrocytes. **(A)** Experimental workflow schematic outlining immunoprecipitation (IP) of CaMKK2 and/or Sox9 from primary WT MACs treated without or with 10 ng/ml IL-1β for 1h or 24 h (1d) followed by immunoblotting. **(B)** Immunoblots (IBs) containing αCaMKK2 or IgG immunoprecipitates probed with antibodies against Ubr4, Psmd2, Sox9 and CaMKK2Black arrows indicate a higher form of Sox9 (>100 kDa) and the expected ∼65 KDa CaMKK2. **(C)** αSox9 IP products resolved on IBs probed with antibodies against Ubr4, Psmd2, Ubiquitin, Sox9 and CaMKK2. Input blots are shown on the left. Black arrows indicate >100 kDa species of ubiquitin, >100 kDa and ∼56 kDa forms of Sox9 and >100 kDa and ∼65 KDa forms of CaMKK2. **(D)** Scheme showing αCaMKK2 and/or αSox9 IP of lysates from WT MACs expressing WT CaMKK2 or its D311A kinase-inactive mutant followed by resolution of bound proteins through immunoblotting. **(E)** IBs of αCaMKK2 or IgG IP of MACs expressing Lentivirus vector only (V), CaMKK2-WT or CaMKK2-D311A probed for Ubr4, Psmd2, Psmc5 (Rpt6), Sox9 and CaMKK2. Black arrows show ∼45 kDa Psmc5, >100 kDa and ∼56 kDa species of Sox9 and ∼68 KDa CaMKK2 doublet. Asterisk in Psmc5 IB indicates IgG heavy chain. **(F)** IBs of αSox9 IP of MACs expressing V, CaMKK2-WT or CaMKK2-D311A probed for Ubr4, Psmd2, ubiquitin, Sox9 and CaMKK2. Black arrows show >100 KDa ubiquitin, >100 kDa (faint band), ∼65 kDa and ∼56 kDa species of Sox9; and >100 kDa and ∼65 KDa (doublet) forms of CaMKK2. **(B-C; E-F)**. Input blots, also probed for b-Actin, are shown; representative IBs from n≥4 biological replicates shown.

### Polyubiquitinated Sox9 complexes with CaMKK2 in naïve and inflamed chondrocytes

Examination of CaMKK2 immunoprecipitates revealed the presence of a higher molecular weight (MW) form of Sox9 (≥ 100 kDa) in IL-1β-treated WT MACs **(Fig. 4B, arrow)**. Notably, no band corresponding to Sox9’s normal MW of 56 kDa was detected in CaMKK2 co-IP. The higher MW Sox9 persisted in CaMKK2 co-IP from MACs treated with IL-1β for 24 h, even though total Sox9 levels were markedly diminished **(Fig. 4B, Input)**. Further, probing Sox9 IP complexes for CaMKK2 revealed two bands: one at its expected MW (∼65 kDa) and the other at ≥100 kDa in both naïve and inflamed MACs **(Fig. 4C, arrow)**. We also observed multiple bands for Sox9, including one at 56 kDa and another at ≥ 100 kDa, among Sox9 immunoprecipitates under both conditions **(Fig. 4C, arrows)**. The higher MW Sox9 bands were retained in Sox9-IP complexes from MACs treated with IL-1β for 24 h, despite a marked reduction in the 56 kDa form and total Sox9 **(Fig. 4C, Input)**. Ubiquitin co-IP’d with Sox9 as a ≥100 kDa conjugate in naïve and IL-1β-treated MACs **(Fig. 4C)**. Thus, ≥100 kDa forms of CaMKK2, Sox9 and ubiquitin co-IP’d with Sox9 in naïve and inflamed MACs, whereas a ≥100 kDa Sox9 species coprecipitated with CaMKK2 in inflamed MACs **(Fig. 4C)**. Interestingly, a 110 kDa polyubiquitinated form of Sox9 in chondrocytes has been previously reported [37]. Together, these findings indicate that CaMKK2 associates with polyubiquitinated Sox9 in chondrocytes and suggest a role for CaMKK2 in regulating proteasome activity and Sox9 turnover.

### CaMKK2 kinase activity is dispensable for its association with UPS subunits and ubiquitinated Sox9

To determine whether CaMKK2 kinase activity is required for its interactions with UPS, we overexpressed kinase-active (WT) CaMKK2 or its kinase-inactive mutant (CaMKK2-D311A) in naïve primary WT MACs using Lentiviruses as before [17], and performed co-IP with αCaMKK2 or αSox9 antibodies **(Fig. 4D)**. CaMKK2 immunoprecipitated Ubr4, Psmd2, and both the 56 kDa and ≥100 kDa forms of Sox9 from MACs expressing kinase-active and/or kinase-inactive CaMKK2 **(Fig. 4E)**. The ≥100 kDa Sox9 species was decreased in CaMKK2-IP from MACs expressing kinase-active CaMKK2, whereas it was enriched and the 56 kDa Sox9 diminished, in IP complexes from MACs expressing kinase-inactive CaMKK2 **(Fig. 4E, arrows)**. Notably, total Sox9 was diminished only when kinase-active CaMKK2, not the kinase-inactive version, was expressed **(Fig. 1D, 2D, and input in 4E**-**4F**).

Analysis of Sox-9 IP complexes revealed interactions with Ubr4, Psmd2, ≥100 kDa ubiquitin and both the 65 kDa and >100 kDa forms of CaMKK2 in MACs overexpressing either WT or kinase-inactive CaMKK2 **(Fig. 4F)**. Interestingly, Sox9 interactions with ≥100 kDa forms of ubiquitin and CaMKK2 were stronger in MACs expressing kinase-inactive CaMKK2, whereas the amount of >100 kDa ubiquitin was markedly lower in Sox9 complexes from MACs expressing kinase-active CaMKK2 **(Fig. 4F)**. Thus, CaMKK2 kinase activity is dispensable for its association with Ubr4, Psmd2 and ubiquitinated Sox9 but it is needed to promote Sox9 turnover.

### CaMKK2 binds to and phosphorylates Psmc5 on Serine 136

Our LC-MS/MS data revealed the presence of the 19S base ATPase subunit Psmc5 (Rpt6) in CaMKK2 IP complexes from IL-1β-treated MACs **(Fig. 3B, 3D)**, and we confirmed these interactions using IP-immunoblotting **(Fig. 4E)**. Interestingly, Psmc5 (∼45 kDa) was more enriched in CaMKK2-immunoprecipitates from MACs expressing empty vector, than those expressing kinase-active CaMKK2, whereas the Psmc5 band was undetectable in CaMKK2 IP complexes from MACs expressing kinase-inactive CaMKK2 **(Fig. 4E, arrow)**, indicating that CaMKK2 kinase activity is required for its interaction with Psmc5 in chondrocytes.

Psmc5 is a phosphoprotein, shown to be directly phosphorylated at Ser120 (S120) by CaMKII and by cAMP-dependent protein kinase (PKA), another Ser/Thr kinase [38, 39]. We therefore investigated whether CaMKK2 phosphorylates Psmc5, by performing in vitro kinase assays using purified recombinant (r) human CAMKK2 and PSMC5 proteins, in the presence of Ca^2+^ and calmodulin **(Fig. 5A)**. CAMKIV, a canonical substrate of CAMKK2, was used as a positive control. The presence of a strong phospho-CAMKIV band (∼ 68 kDa) confirmed that the rCAMKK2 is an active kinase **(Fig. 5B)**. We also observed a band around ∼45 kDa (MW of PSMC5) that is immunoreactive to the pan-phospho Ser/Thr antibody in kinase reactions containing rCAMKK2 and rPSMC5, whereas this band was absent in reactions lacking the kinase **(Fig. 5B)**. These data indicate that CAMKK2 directly phosphorylates PSMC5. To identify the CAMKK2 phosphosite(s) on PSMC5, we performed LC-MS/MS analysis of tryptic digested kinase reactions **(Fig. 5A)** and identified a phosphopeptide ILPNKVDPLVsLMMVEK (126-142) containing phospho-Ser136 (pS136) **(Fig. 5C, Supplementary Fig. 3A-B)**. In two independent experiments, PSMC5 pS136 containing phosphopeptides were identified by 6-14 peptide-spectrum matches (PSMs) in kinase reactions containing CAMKK2 and PSMC5 but were absent in reactions lacking CAMKK2 **(Fig. 5C, Supplementary Fig. 4-8)**. S136 phosphorylation of PSMC5 was verified with a characteristic neutral loss of a phosphoric acid and by the characteristic b and y ion sequences as shown in the representative fragmentation spectra in **Fig. 5C**. The PSMC5 pS136 site was previously identified in a quantitative phosphoproteomics study as an evolutionarily conserved phosphorylation event during the human cell cycle, suggesting a role in mitosis [40]. Given the interaction of CaMKK2 with UPS and the requirement of CaMKK2 kinase activity in Sox9 turnover, we surmised that PSMC5-S136 phosphorylation by CAMKK2 stabilizes UPS and/or enhances proteasome function in chondrocytes.

**Figure 5:**
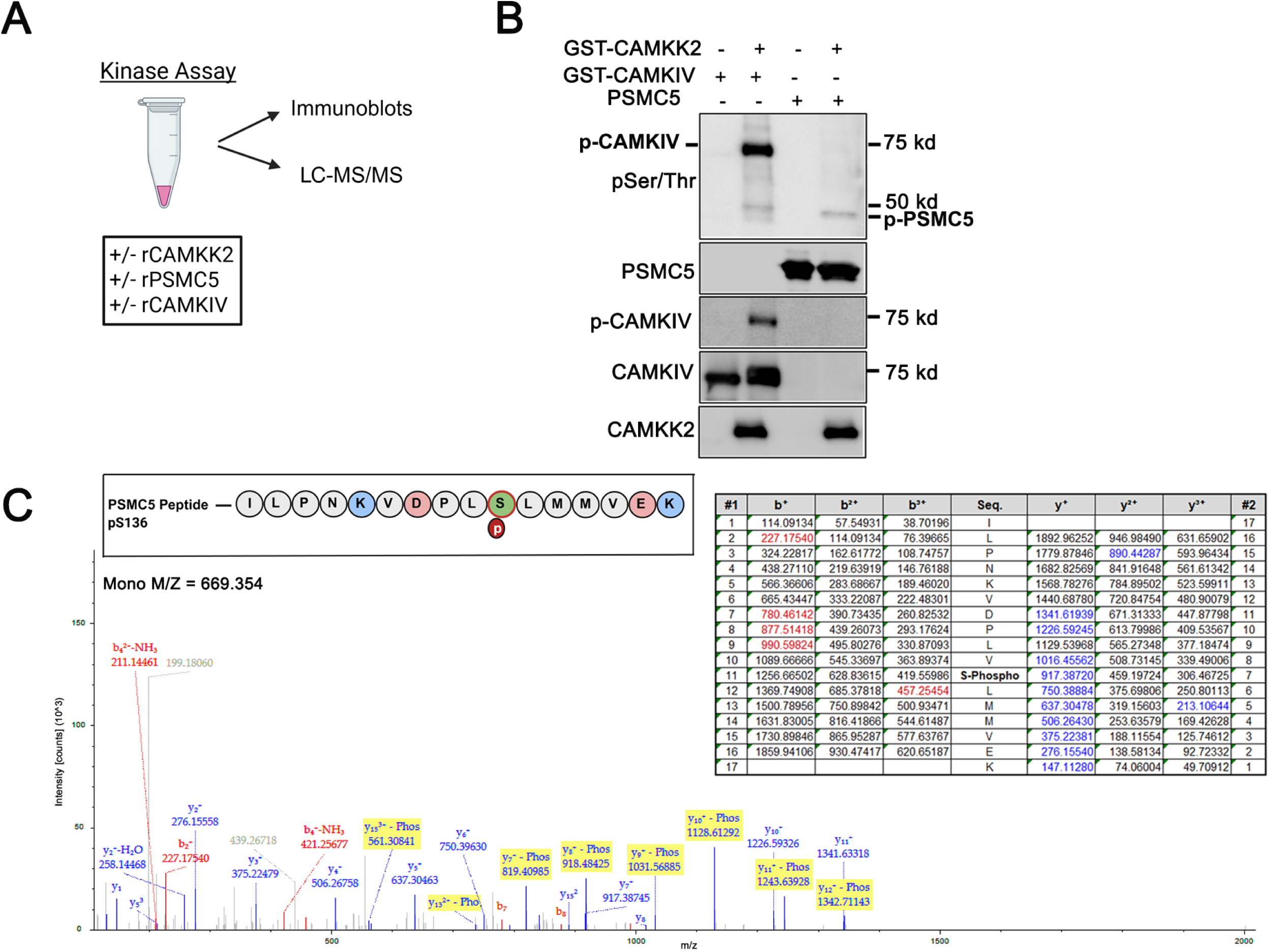
Direct phosphorylation of Psmc5/Rpt6 at Ser136 by CaMKK2. **(A)** Schematic outline of kinase assays with or without recombinant (r) human CAMKK2 in reactions containing human rPSMC5 or rCAMKIV as substrates. The kinase assays were then resolved on IBs or by LC-MS/MS to identify potential phosphosite(s). **(B)** IBs of kinase assay reactions with or without GST-rCAMKK2, GST-rCAMKIV or His-tagged rPSMC5 probed for immunoreactivity to antibodies against pan phospho (p)-Ser/Thr, pS120-PSMC5, pT196/200-CAMKIV as well as total PSMC5, CAMKIV, and CAMKK2. Bands corresponding to pPSMC5 and pCaMKIV are indicated in the pan-Ser/Thr IB. **(C)** A representative LC-MS/MS fragmentation spectrum of phosphopeptide ILPNKVDPL**S**LMMVEK containing the novel Ser136 phosphorylation site on PSMC5 in kinase reactions containing rCAMKK2. Graph depicting the m/z spectra of intensity counts and a table summarizing the characteristic b and y ion series and indicating the neutral loss of a phosphoric acid for S136 in the phosphopeptide ILPNKVDPL**S**LMMVEK are shown. Additional phospho-S136 PSM data (LC-MS/MS) from two independent biological replicates are shown in supplemental data

### CaMKK2 kinase activity is required for its upregulation of chondrocyte proteasome activity

We next measured the proteasome activity in primary WT and *Camkk2^-/-^*MACs, untreated or treated with IL-1β (10 ng/mL, 24 h), using a fluorogenic peptide substrate which releases fluorescent moiety upon cleavage by the active 20S proteasome **(Fig. 6A)**. Fluorescence intensity at ∼480 nm reflects proteolytic activity and enables quantitative assessment of proteasome function. Basal proteasome activity in naïve *Camkk2^-/-^* MACs was 20% lower than WT **(Fig. 6B)**. Whereas IL-1β treatment increased proteasome activity in both genotypes, *Camkk2^-/-^* MACs treated with IL-1β exhibited significantly lower proteasome activity than IL-1β-treated WT MACs (**Fig. 6B)**. Further, expression kinase-active CaMKK2 enhanced total proteasome function by 20% in primary human chondrocytes and by 13% in primary MACs, whereas kinase-inactive CaMKK2 had no effect **(Fig. 5C, 5D)**. These findings establish that CaMKK2 kinase activity is essential for phosphorylating Psmc5 on S136, enhancing proteasome function and promoting Sox9 degradation in naïve and inflamed chondrocytes.

**Figure 6:**
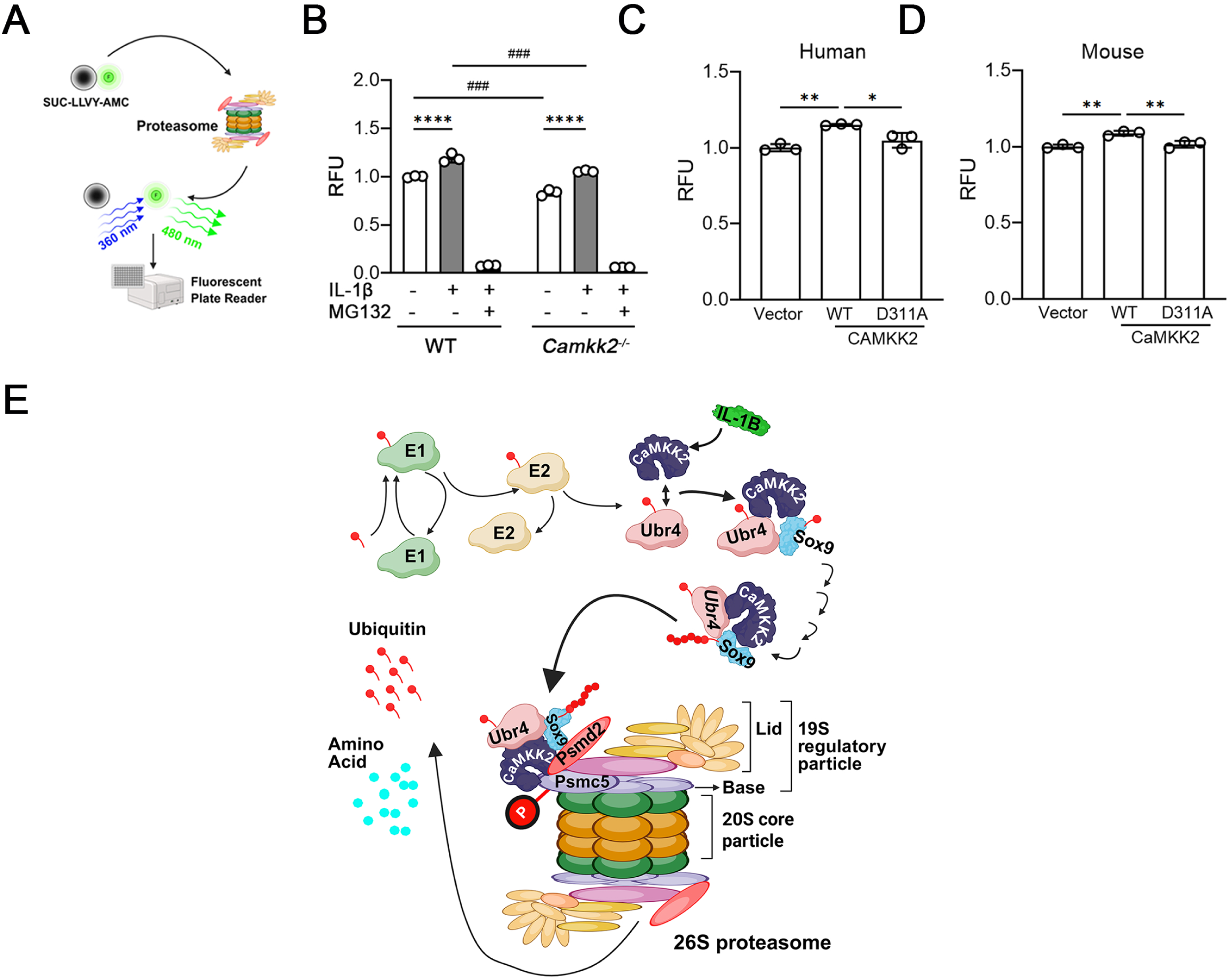
Chondrocyte proteasome activity is regulated by kinase-active CaMKK2. **(A)** Scheme outlining the quantitative assessment of proteasome function from cell lysates using a fluorogenic peptide substrate SUC-LLVY-AMC of the 26S proteasome. Cleavage of the peptide by the 20S proteasome releases AMC-480 nm fluorescent fragment, and the 480 nm fluorescence intensity is quantified using a plate reader. **(B)** Graph showing relative fluorescence units (RFU) indicating 480 nm fluorescence intensity in WT and *Camkk2^-/-^* primary MACs treated with or without 10 ng/ml IL-1β for 24 h, in the presence or absence of the proteasome inhibitor MG132. N=3; Error bars = SD; * indicates *p*-values for within genotype comparison and # for WT vs. *Camkk2^-/-^*comparisons; \*\*\*\**p* < 0.0001; ^###^ *p*<0.001. **(C-D)** RFU (480 nm fluorescence intensity) in primary healthy human chondrocytes **(C)** and/or WT MACs **(D)** expressing Lentivirus vector alone, WT-CaMKK2 or D311A mutant. N=3; SD; * *p*<0.05; ** *p* < 0.01. **(E)** Working model of the mechanism of regulation of chondrocyte UPS and Sox9 levels by CaMKK2. In chondrocytes, Ubr4, a massive 600 kDa E3/E4 ligase that extends ubiquitin chains on pre-ubiquitinated substrates, binds to CaMKK2, which is stimulated by IL-1β. The CaMKK2-Ubr4 complex binds to monoubiquitinated substrates such as Sox9, and Ubr4 along with its co-factor Kcmf1 (not pictured) elongates the ubiquitin chains on Sox9. Ubr4-polyubiquitinated Sox9 bound CaMKK2 then binds to seven Psmd proteins that make up the lid subcomplex and all six AAA ATPase proteins that constitute the base subcomplex of the 19S proteasome RP. Additionally, CaMKK2 phosphorylates Psmc5 on S136, a phosphosite previously identified to be associated with cell mitosis. We hypothesize that through these interactions with Ubr4 and 19S proteasome, and by directly phosphorylating Psmc5, CaMKK2 functionally stabilizes the 26S proteasome, as proteasome activity is attenuated in naïve and inflamed *Camkk2^-/-^* MACs, compared to WT. The kinase activity of CaMKK2 is critical to its regulation of proteasome function as the kinase-inactive CaMKK2 mutant was unable to stimulate proteasome activity.

## Discussion

This study uncovers a previously unrecognized role for CaMKK2 in chondrocyte proteostasis and identifies Psmc5 as a novel CaMKK2 substrate. Loss or inhibition of CaMKK2 elevates Sox9 in articular chondrocytes and protects it from inflammatory cue–induced degradation. Mechanistically, CaMKK2 complexes with E3 ubiquitin ligase Ubr4, multiple 19S RP subunits, and polyubiquitinated Sox9, thereby linking inflammatory signaling to UPS-mediated Sox9 turnover. CaMKK2 also interacts with and phosphorylates the 19S RP AAA ATPase Psmc5 at S136, and its kinase function is required for enhanced chondrocyte proteasome activity and Sox9 degradation **(Fig. 6E)**. Consistent with this model, Sox9 levels are increased in naïve *Camkk2^-/-^* cartilage and preserved after DMM surgery or pharmacologic CaMKK2 inhibition in murine and human OA samples, where CaMKK2 is elevated and Sox9 is reduced.

Ubr4, the most enriched protein in the CaMKK2-IP complex, is a 600 kDa multifunctional E3/E4 ubiquitin ligase that is central to protein quality control in multiple cell types[35, 41–44]. Functionally, Ubr4 extends polyubiquitin chains on substrates for proteasomal degradation [35]. The interaction of CaMKK2 with Ubr4 suggests a mechanism whereby CaMKK2 may scaffold the ubiquitination machinery around target proteins such as Sox9, facilitating their polyubiquitination and subsequent proteasomal targeting. CaMKK2 also associates with two Ubr4 cofactors: Kcmf2, another E3 ligase, and calmodulin, the Ca^2+^-binding protein that regulates CaMKK2 activity [35, 45]. We previously showed CaMKK2 to mediate IL-1β-induced activation of IL-6-STAT3-MMP13 pathway in chondrocytes [17, 18]. Inflammatory cytokines such as IL-1β trigger transient increases in intracellular Ca^2+^, which binds to calmodulin to activate CaMKK2. Taken together, our findings underscore the integration of Ca^2+^ signaling with UPS function through CaMKK2.

CaMKK2 also associates with multiple 19S RP subunits which recognize polyubiquitinated substrates and mediate their deubiquitination, unfolding, and translocation into the 20S catalytic core for degradation [36, 46]. Interestingly, CaMKK2 kinase activity was not necessary for its binding to Ubr4, Psmd2 and polyubiquitinated Sox9, but it was required for its regulation of the chondrocyte proteasome activity and for its phosphorylation of Psmc5 on Ser136. Psmc5 forms part of the six 19S RP hexameric AAA ATPase ring (Psmc1–6) base, which provides the ATP-dependent mechanical force for substrate deubiquitination and translocation into the 20S catalytic chamber [46]. Phosphorylation of Psmc5 at S136 by CaMKK2 is therefore poised to modulate proteasome efficiency.

Limitations of our study include our reliance on a single commercially available antibody (rabbit anti-CaMKK2-NT antibody; 033168; US Biological, Salem, MA) for specifically immunoprecipitating murine CaMKK2 [17, 18, 47], which partly hindered our efforts to validate CaMKK2 interactions with additional 19S RP subunits identified by proteomics. Moreover, the availability of high-quality antibodies that recognize phosphorylated and unmodified forms of the interacting proteins was another impediment. Future studies using recombinant proteins may identify additional CaMKK2 substrates within the UPS and determine how their phosphorylation by CaMKK2 regulates the proteasome activity in chondrocytes.

In conclusion, CaMKK2 integrates inflammatory and Ca²⁺ signaling with UPS-mediated proteostasis in chondrocytes through dual, complementary mechanisms: a scaffolding function that recruits ubiquitination proteasome machinery to polyubiquitinated substrates like Sox9, and a catalytic function that phosphorylates Psmc5 and enhances proteasome activity. Through these coordinated roles, CaMKK2 emerges as a key regulator of chondrocyte protein homeostasis and cartilage integrity under inflammatory stress, positioning it as a promising therapeutic target against OA.

## Acknowledgments

The authors thank Dr. Lilian Plotkin, Drew M. Brown and staff at the Musculoskeletal Histology Core Facility of the ICMH Core of the Indiana Clinical Translational Sciences Institute (CTSI) for help with histology. LC-MS/MS was performed by IUSM Center for Proteome Analysis.

## Data Availability

Raw and processed mass spectrometry data are uploaded to ProteomeXchange partner MassIVE and password protected.

## Author Contributions

XD designed and performed experiments, analyzed data, prepared figures, and wrote the manuscript; YL performed experiments and analyzed data; ESY provided reagents, contributed to experimental design and data interpretation, and edited the manuscript; KH and EHD performed and analyzed LC-MS/MS assays, prepared figures, and wrote methods; EHD and ALM interpreted proteomics data and edited the manuscript; and US conceived and supervised the study, secured funding, analyzed data, prepared figures, and drafted the manuscript.

## References

1. Global, regional, and national burden of osteoarthritis, 1990-2020 and projections to 2050: a systematic analysis for the Global Burden of Disease Study 2021. Lancet Rheumatol, 2023. 5(9): p. e508–e522.

2. Dilley, J.E., et al., Post-traumatic osteoarthritis: A review of pathogenic mechanisms and novel targets for mitigation. Bone Rep, 2023. 18: p. 101658.

3. Pritchard, S. and F. Guilak, Effects of interleukin-1 on calcium signaling and the increase of filamentous actin in isolated and in situ articular chondrocytes. Arthritis Rheum, 2006. 54(7): p. 2164–74.

4. Lee, W., et al., Inflammatory signaling sensitizes Piezo1 mechanotransduction in articular chondrocytes as a pathogenic feed-forward mechanism in osteoarthritis. Proc Natl Acad Sci U S A, 2021. 118(13).

5. Riggs, K.C. and U. Sankar, Inflammatory mechanisms in post-traumatic osteoarthritis: a role for CaMKK2. Immunometabolism (Cobham), 2023. 5(4): p. e00031.

6. Kaokhum, N., et al., The Mechano-Ubiquitinome of Articular Cartilage: Differential Ubiquitination and Activation of a Group of ER-Associated DUBs and ER Stress Regulators. Mol Cell Proteomics, 2022. 21(12): p. 100419.

7. Liu-Bryan, R. and R. Terkeltaub, Emerging regulators of the inflammatory process in osteoarthritis. Nature Reviews Rheumatology, 2015. 11(1): p. 35–44.

8. Olivotto, E., et al., Pathophysiology of osteoarthritis: canonical NF-κB/IKKβ-dependent and kinase-independent effects of IKKα in cartilage degradation and chondrocyte differentiation. RMD Open, 2015. 1(Suppl 1): p. e000061.

9. Lee, D. and J.H. Hong, Physiological Overview of the Potential Link between the UPS and Ca(2+) Signaling. Antioxidants (Basel), 2022. 11(5).

10. Haseeb, A., et al., SOX9 keeps growth plates and articular cartilage healthy by inhibiting chondrocyte dedifferentiation/osteoblastic redifferentiation. Proc Natl Acad Sci U S A, 2021. 118(8).

11. Akiyama, H., et al., The transcription factor Sox9 is degraded by the ubiquitin–proteasome system and stabilized by a mutation in a ubiquitin-target site. Matrix Biology, 2005. 23(8): p. 499–505.

12. Mei, Z., et al., Chondrocyte fatty acid oxidation drives osteoarthritis via SOX9 degradation and epigenetic regulation. Nature Communications, 2025. 16(1): p. 4892.

13. Hu, W., et al., Bortezomib prevents the expression of MMP-13 and the degradation of collagen type 2 in human chondrocytes. Biochemical and Biophysical Research Communications, 2014. 452(3): p. 526–530.

14. Quan, R., et al., Effects of a proteasome inhibitor on the NF-κB signalling pathway in experimental osteoarthritis. Scandinavian Journal of Rheumatology, 2013. 42(5): p. 400–407.

15. Wang, W., et al., Attenuated Joint Tissue Damage Associated With Improved Synovial Lymphatic Function Following Treatment With Bortezomib in a Mouse Model of Experimental Posttraumatic Osteoarthritis. Arthritis Rheumatol, 2019. 71(2): p. 244–257.

16. Radwan, M., et al., Protection against murine osteoarthritis by inhibition of the 26S proteasome and lysine-48 linked ubiquitination. Annals of the Rheumatic Diseases, 2015. 74(8): p. 1580–1587.

17. Dilley, J.E., et al., CAMKK2 is upregulated in primary human osteoarthritis and its inhibition protects against chondrocyte apoptosis. Osteoarthritis Cartilage, 2023.

18. Mevel, E., et al., Systemic inhibition or global deletion of CaMKK2 protects against post-traumatic osteoarthritis. Osteoarthritis Cartilage, 2022. 30(1): p. 124–136.

19. Lefebvre, V., M. Angelozzi, and A. Haseeb, SOX9 in cartilage development and disease. Curr Opin Cell Biol, 2019. 61: p. 39–47.

20. Kellgren, J.H. and J.S. Lawrence, Radiological assessment of osteo-arthrosis. Ann Rheum Dis, 1957. 16(4): p. 494–502.

21. Mosley, A.L., et al., Highly reproducible label free quantitative proteomic analysis of RNA polymerase complexes. Mol Cell Proteomics, 2011. 10(2): p. M110 000687.

22. Grecco, G.G., et al., A multi-omic analysis of the dorsal striatum in an animal model of divergent genetic risk for alcohol use disorder. J Neurochem, 2021. 157(4): p. 1013–1031.

23. Orsburn, B.C., Proteome Discoverer—A Community Enhanced Data Processing Suite for Protein Informatics. Proteomes, 2021. 9(1): p. 15.

24. Bell, D.M., et al., SOX9 directly regulates the type-II collagen gene. Nat Genet, 1997. 16(2): p. 174–8.

25. Ng, L.J., et al., SOX9 binds DNA, activates transcription, and coexpresses with type II collagen during chondrogenesis in the mouse. Dev Biol, 1997. 183(1): p. 108–21.

26. Sekiya, I., et al., SOX9 Enhances Aggrecan Gene Promoter/Enhancer Activity and Is Up-regulated by Retinoic Acid in a Cartilage-derived Cell Line, TC6*. Journal of Biological Chemistry, 2000. 275(15): p. 10738–10744.

27. Ouyang, Y., et al., Overexpression of SOX9 alleviates the progression of human osteoarthritis in vitro and in vivo. Drug Des Devel Ther, 2019. 13: p. 2833–2842.

28. Fukui, N., et al., Regional differences in chondrocyte metabolism in osteoarthritis: A detailed analysis by laser capture microdissection. Arthritis & Rheumatism, 2008. 58(1): p. 154–163.

29. Tokumitsu, H., et al., STO-609, a specific inhibitor of the Ca(2+)/calmodulin-dependent protein kinase kinase. J Biol Chem, 2002. 277(18): p. 15813–8.

30. Kim, S., et al., Tankyrase inhibition preserves osteoarthritic cartilage by coordinating cartilage matrix anabolism via effects on SOX9 PARylation. Nature Communications, 2019. 10(1): p. 4898.

31. Lee, D.H. and A.L. Goldberg, Proteasome inhibitors: valuable new tools for cell biologists. Trends Cell Biol, 1998. 8(10): p. 397–403.

32. Hunt, L.C., et al., A Key Role for the Ubiquitin Ligase UBR4 in Myofiber Hypertrophy in Drosophila and Mice. Cell Reports, 2019. 28(5): p. 1268–1281.e6.

33. Parsons, K., Y. Nakatani, and M.D. Nguyen, p600/UBR4 in the central nervous system. Cell Mol Life Sci, 2015. 72(6): p. 1149–60.

34. Nakatani, Y., et al., p600, a unique protein required for membrane morphogenesis and cell survival. Proceedings of the National Academy of Sciences, 2005. 102(42): p. 15093–15098.

35. Grabarczyk, D.B., et al., Architecture of the UBR4 complex, a giant E4 ligase central to eukaryotic protein quality control. Science, 2025. 389(6763): p. 909–914.

36. Budenholzer, L., et al., Proteasome Structure and Assembly. J Mol Biol, 2017. 429(22): p. 3500–3524.

37. Hattori, T., et al., E6-AP/UBE3A protein acts as a ubiquitin ligase toward SOX9 protein. J Biol Chem, 2013. 288(49): p. 35138–48.

38. Djakovic, S.N., et al., Regulation of the Proteasome by Neuronal Activity and Calcium/Calmodulin-dependent Protein Kinase II*. Journal of Biological Chemistry, 2009. 284(39): p. 26655–26665.

39. Zhang, F., et al., Proteasome function is regulated by cyclic AMP-dependent protein kinase through phosphorylation of Rpt6. Journal of Biological Chemistry, 2007. 282(31): p. 22460–22471.

40. Olsen, J.V., et al., Quantitative phosphoproteomics reveals widespread full phosphorylation site occupancy during mitosis. Sci Signal, 2010. 3(104): p. ra3.

41. Hunt, L.C., et al., Antagonistic control of myofiber size and muscle protein quality control by the ubiquitin ligase UBR4 during aging. Nat Commun, 2021. 12(1): p. 1418.

42. Kim, S.T., et al., The N-recognin UBR4 of the N-end rule pathway is required for neurogenesis and homeostasis of cell surface proteins. PLoS One, 2018. 13(8): p. e0202260.

43. Haakonsen, D.L., et al., Stress response silencing by an E3 ligase mutated in neurodegeneration. Nature, 2024. 626(8000): p. 874–880.

44. Jeong, D., et al., Tumor-promoting UBR4 coordinates impaired mitophagy–associated senescence and lung adenocarcinoma pathogenesis. Proceedings of the National Academy of Sciences, 2025. 122(25): p. e2425015122.

45. Means, A.R., The Year in Basic Science: calmodulin kinase cascades. Molecular endocrinology, 2008. 22(12): p. 2759–65.

46. Tanaka, K., The proteasome: overview of structure and functions. Proc Jpn Acad Ser B Phys Biol Sci, 2009. 85(1): p. 12–36.

47. Williams, J.N., et al., Osteocyte-Derived CaMKK2 Regulates Osteoclasts and Bone Mass in a Sex-Dependent Manner through Secreted Calpastatin. Int J Mol Sci, 2023. 24(5).

