## Supplementary Figures 1-8 for "A CAMKK2-UBR4-19S Proteasome Axis Regulates Chondrocyte Proteostasis and SOX9 Stability"

^1^Department of Anatomy, Cell Biology and Physiology, Indiana University School of Medicine, Indianapolis, IN 46202, USA; ^2^Indiana Center for Musculoskeletal Health, Indiana University School of Medicine, Indianapolis, IN 46202, USA; ^3^Department of Biochemistry, Molecular Biology and Pharmacology; ^4^Center for Proteome Analysis; ^5^Center for Computational Biology and Bioinformatics, Indiana University School of Medicine, Indianapolis, IN 46202, USA.

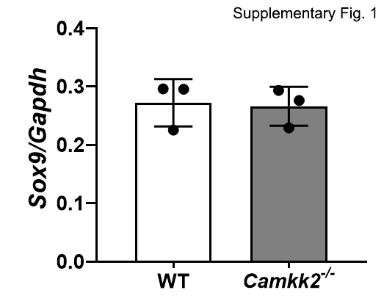

**Supplementary Figure 1: Loss of CaMKK2 does not affect *Sox9* mRNA levels in MACs.** *Sox9* mRNA normalized to *Gapdh* in naïve primary WT and *Camkk2^-/-^* MACs indicating no differences. N=3 biological replicates.

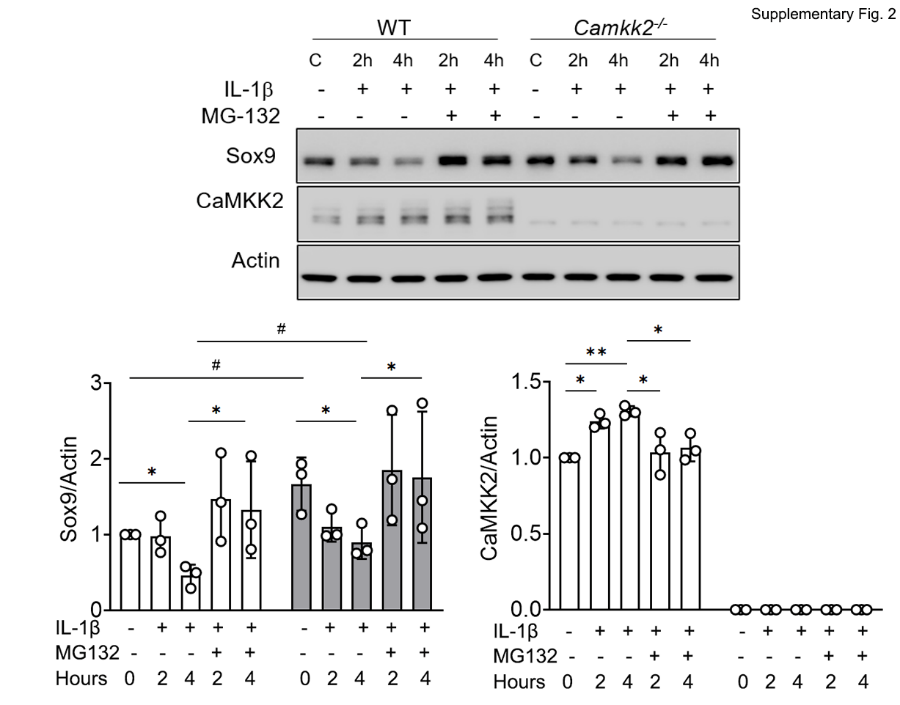

**Supplementary Figure 2: IL-1β-mediated Sox9 degradation occurs via UPS in both WT and *Camkk2^-/-^* MACs.** Immunoblots probed for Sox9, CaMKK2 and b-Actin and quantification. Cell lysates were prepared from primary WT and *Camkk2^-/-^* MACs treated with 10 ng/ml of IL-1β with or without 1 µM MG-132 for indicated timepoints. N=3; SD; * indicates *p*-values for within genotype comparison and # for WT vs. *Camkk2^-/-^* comparisons; ^*^ *p*<0.05; ** *p* < 0.01 and ^#^ *p*<0.05; ^##^ *p*<0.01.

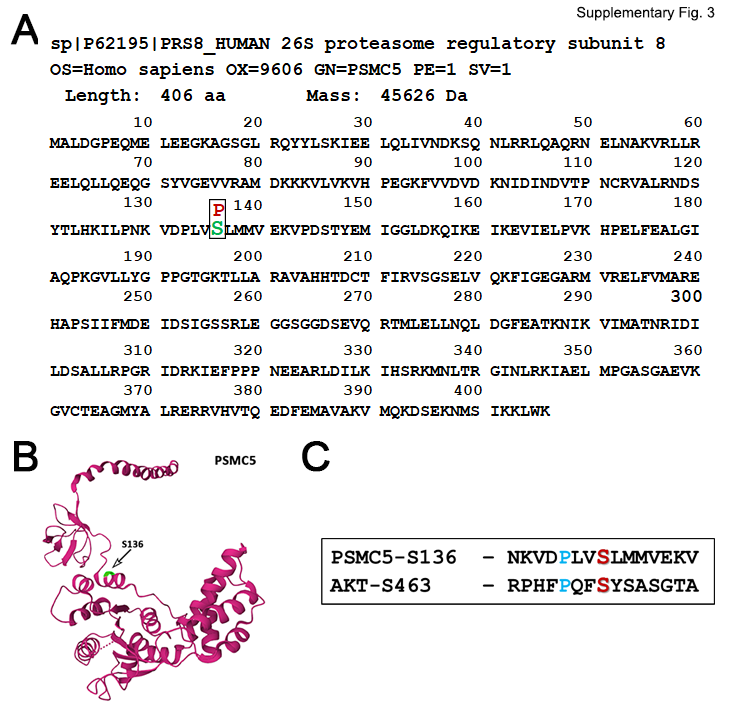

**Supplementary Figure 3: Identification of a novel pS136 on PSMC5 phosphorylated by CaMKK2. (A)** Human PSMC5 (P62195) protein sequence showing the S136 phosphosite identified based on LC-MS/MS data (Serine – Green; phospho (P) site – Red). **(B)** PSMC5 3D structure from Protein Data Bank (PDK), with S136 highlighted in Green. **(C)** PSMC5 and AKT phosphopeptides containing the respective CAMKK2 phosphosites – S136 and S463, indicating Proline residue at -3 position.

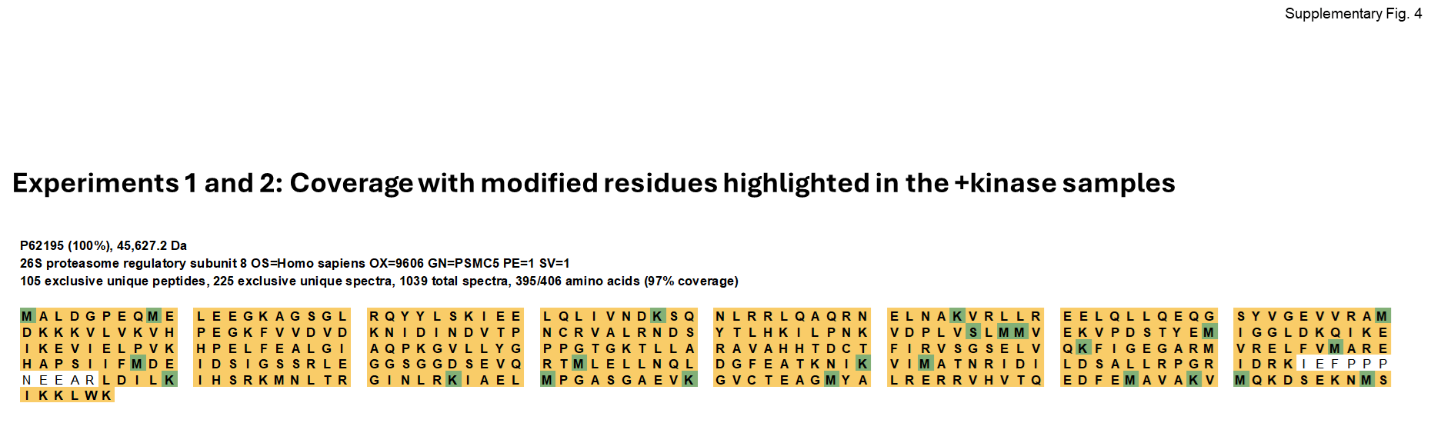

**Supplementary Figure 4. LC-MS/MS raw data 1/5: Coverage with modified residues in PSMC5 in kinase assays containing CaMKK2**. PSMC5 sequences with modified residues highlighted in Green from experiments 1 and 2 in samples containing CaMKK2.

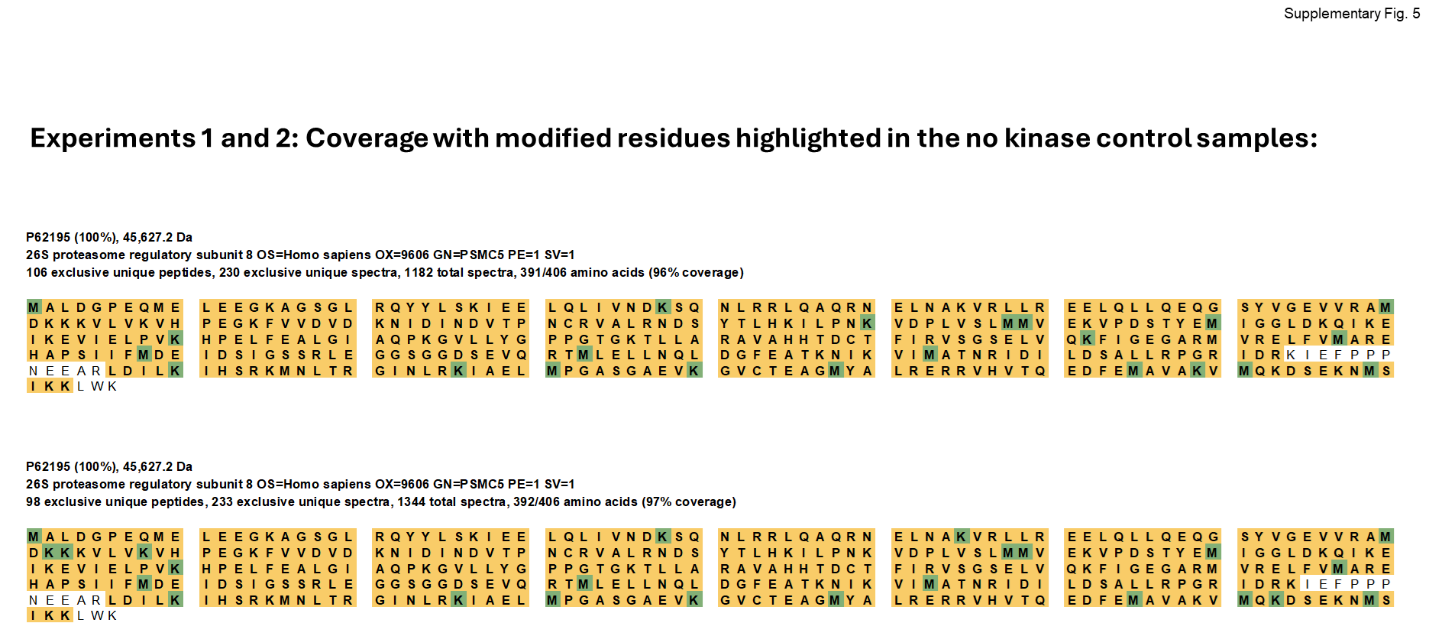

**Supplementary Figure 5. LC-MS/MS raw data 2/5: Coverage with modified residues in PSMC5 in kinase assays lacking CaMKK2**. PSMC5 sequences with modified residues highlighted in Green from experiments 1 and 2 in samples without CaMKK2.

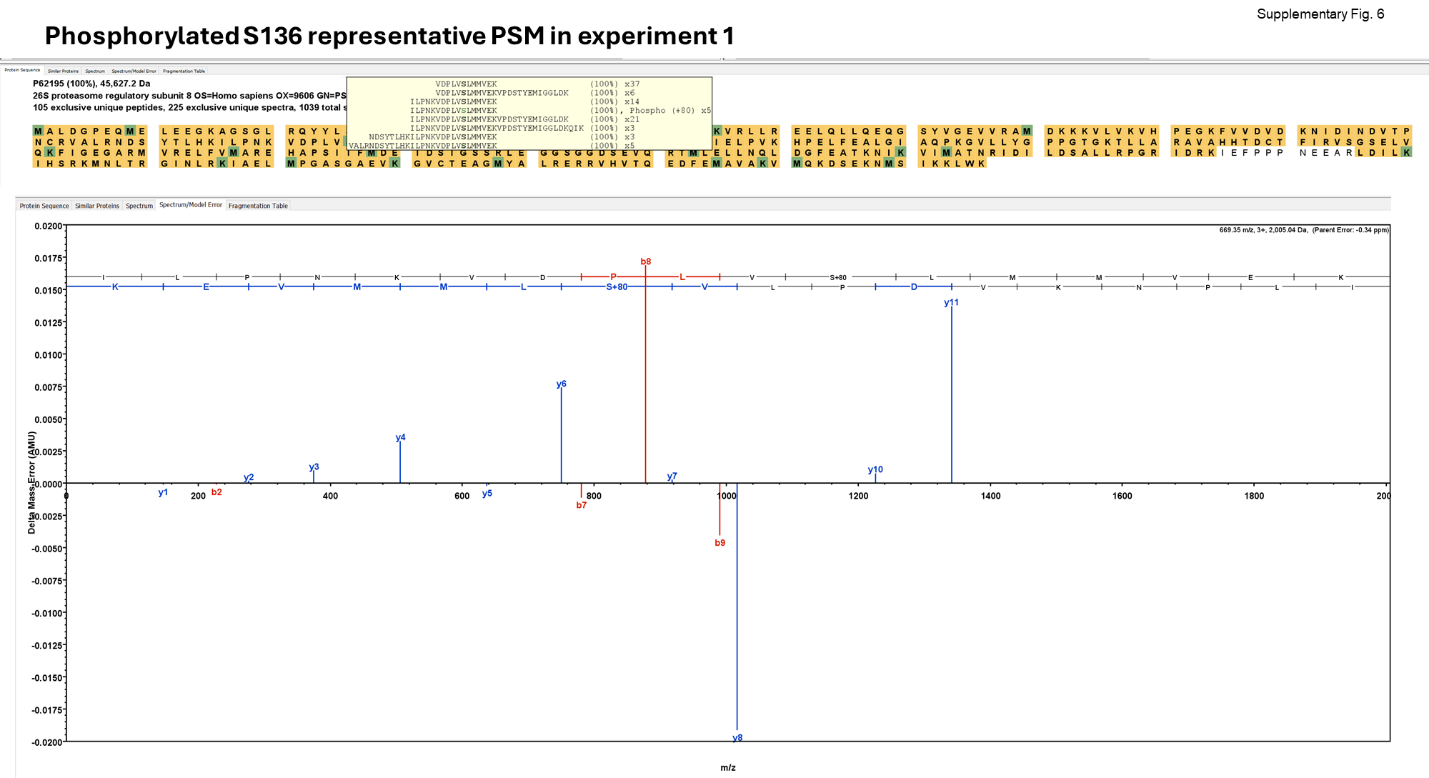

**Supplementary Figure 6. LC-MS/MS raw data 3/5: Phosphorylated PSMC5 S136 PSM from experiment 1 in kinase assays containing CaMKK2**. PSM spectra from experiment 1 in samples containing CaMKK2.

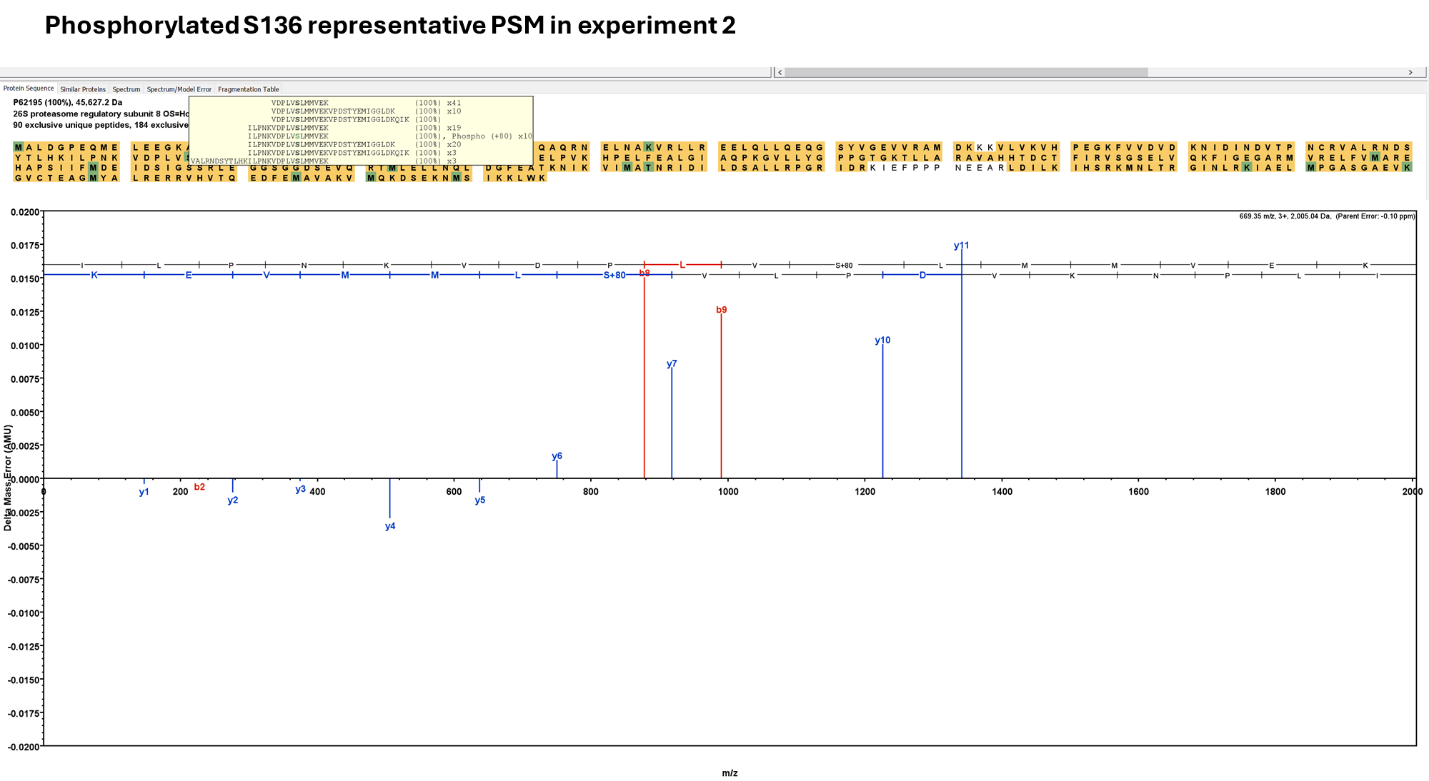

**Supplementary Figure 7. LC-MS/MS raw data 4/5: Phosphorylated PSMC5 S136 PSM from experiment 2 in kinase assays containing CaMKK2**. PSM spectra from experiment 2 in samples containing CaMKK2.

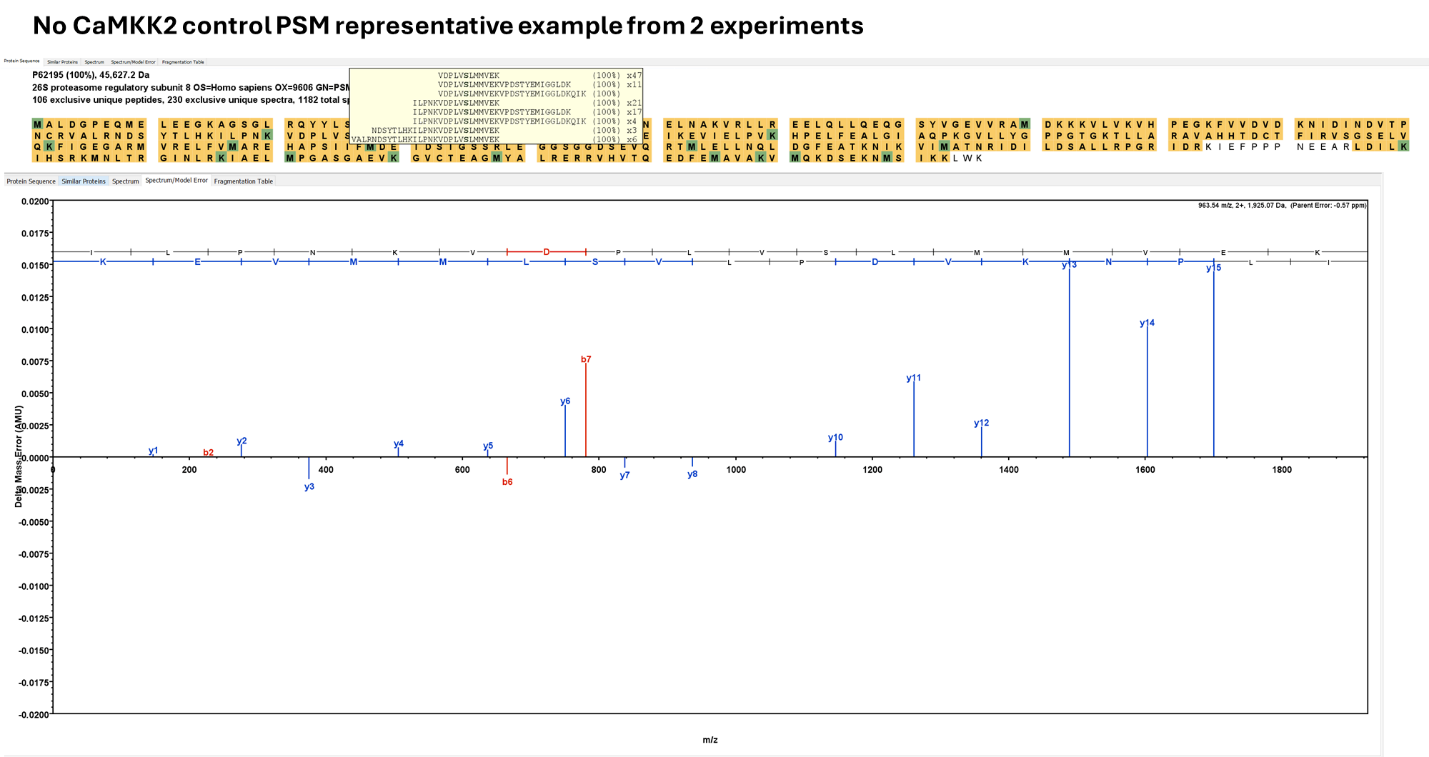
**Supplementary Figure 8. LC-MS/MS raw data 5/5: Representative PSM from n=2 experiments of kinase assays without CaMKK2**. PSM spectra from 2 kinase assays without CaMKK2.
